# Monocyte reprogramming by nociceptors impedes T cell-mediated tumor immunity

**DOI:** 10.64898/2026.01.15.699724

**Authors:** Pavel Hanč, Adrian Blair, Harold R. Neely, Changwei Peng, Irina B. Mazo, Matthias Mack, Ulrich H. von Andrian

## Abstract

Nociceptors – peripheral nervous system neurons that trigger the sensation of pain or itch in response to noxious stimuli – can communicate with leukocytes and modulate immune responses in a context-dependent manner^1,2^. Here, we demonstrate that nociceptors rapidly induce monocytes to adopt an immunosuppressive phenotype, reminiscent of myeloid-derived suppressor cells (MDSCs), which have been linked to impaired anti-tumor immunity. MDSCs are abundant in human bladder cancer, a malignancy where survival is negatively correlated with expression of pro-neurogenic cytokines. Accordingly, in an orthotopic mouse model of urinary bladder carcinoma, incipient tumors promoted dense innervation by nociceptors and infiltration by monocytes that differentiated rapidly into MDSCs. Nociceptors did not affect the development of tumor-specific T cells in draining lymph nodes, but they were essential during the early stages of tumor formation to establish an immunosuppressive tumor microenvironment. Ablation or chemogenetic inhibition of nociceptors prevented the formation of immunosuppressive MDSCs and markedly increased the rate of bladder tumor rejection by T cells. Conversely, monocyte depletion in nociceptor-sufficient mice enabled tumor rejection. Collectively, these findings demonstrate that urothelial tumors co-opt an immunosuppressive neuro-immune axis to evade anti-tumor immunity and identify nociceptors as potential targets for bladder cancer treatment.

## Introduction

The tumor microenvironment (TME) is a complex cellular ecosystem comprising malignant tumor cells, tumor-infiltrating leukocytes, and various stromal components, including fibroblasts, vascular endothelial cells, and nerve fibers^3^. Much recent work has focused on understanding the varied roles of immune cells in the TME and how the balance between pro- and anti-inflammatory processes affects tumorigenesis. Across tumor types, tumor cell killing by tumor antigen-specific CD8^+^ cytotoxic T-cells (CTLs) is considered the primary immunological mechanism to prevent or delay cancer progression^4^. However, CTL activity is frequently compromised, as the TME represents an immunosuppressive milieu that inhibits the activity of tumoricidal cells through a variety of mechanisms. These include the expression of checkpoint molecules such as PD-L1, which directly block the activity of CTLs^5^, as well as the accumulation of immunosuppressive immune cell types, including regulatory T-cells and various myeloid leukocytes. Tumor-associated immunosuppressive myeloid cells can be broadly categorized as tumor-associated macrophages (TAMs) and myeloid-derived suppressor cells (MDSCs)^6^. Based on their phenotype and origin, MDSCs themselves are further divided into Ly6C^+^Ly6G^-^monocytic MDSCs (M-MDSCs) and Ly6C^Lo^Ly6G^+^ polymorphonuclear MDSCs (PMN-MDSCs)^7^.

In contrast to the immune system, the role of various stroma components in the pathobiology of solid tumors is still poorly understood. In particular, the relevance of tumor innervation has only recently begun to receive some attention. For example, recent work has shown that a high density of intratumoral nerve fibers correlates with poor prognosis across multiple tumor types,^8^ suggesting a possible pro-tumorigenic function. The peripheral nervous system comprises the autonomic branch, which regulates most involuntary processes and the somatic branch, comprising motor neurons that control voluntary movements and somato-sensory fibers that transmit sensory information. The cell bodies of somatosensory neurons mainly reside in the dorsal root and trigeminal ganglia from where they project axons toward the periphery. A large population within the somato-sensory system are the nociceptors, a specialized subset of sensory neurons that elicit the sensation of pain or itch in response to noxious stimuli.

Nociceptors are increasingly recognized as key regulators of immune responses; their immunological effects are context-dependent and can be both pro- or anti-inflammatory^1,2^. For example, nociceptors can promote immune responses by communicating with dendritic cells (DCs)^9–11^ and B-cells^12^, but they can also impair immunity against bacterial infections^13–15^ and promote tissue repair^16^ by inhibiting histotoxic actions of neutrophils and macrophages.

Recent work has begun to uncover the importance of nociceptors within the TME^17^. For example, in melanoma, nociceptors impair the formation of non-classical tertiary lymphoid structures^18^ and secrete calcitonin gene-related peptide (CGRP), a neuropeptide that dampens anti-tumor immunity by inducing an exhausted phenotype in tumor-infiltrating CTLs^19^. Analogous observations were also made in head and neck squamous cell carcinoma, where nociceptors promoted CTL exhaustion^20–22^ and induced an immunosuppressive phenotype in bone marrow leukocytes^22^. Conversely, in gastric cancer, nociceptor-derived CGRP promotes tumor growth by acting directly on the tumor cells^23^, while another nociceptive neuropeptide, substance P, can promote metastasis in breast cancer^24^. It is still unknown whether and by what mechanism(s) nociceptors contribute to the pathobiology of other tumor types.

Seeking an answer to this question, we identified human bladder cancer as a candidate tumor type in which high expression of neurotrophic cytokines was negatively correlated with patient survival. Indeed, in a mouse model of orthotopic bladder cancer, ablation of nociceptors dramatically decreased tumor formation. Mechanistically, our studies indicate that incipient tumors stimulated local nociceptive innervation and, simultaneously, promoted the recruitment of migratory monocytes. Within the TME, monocytes that encountered nociceptors were driven to differentiate into M-MDSCs that protected the tumor against CTL-mediated rejection.

Consequently, ablation of either nociceptors or monocytes was sufficient to prevent bladder cancer formation. These results identify an additional, previously unrecognized cellular mechanism by which nociceptors can promote tumor growth and illustrate that nociceptors can affect tumor formation by a variety of cellular and molecular mechanisms that may depend on tumor type and anatomic context.

## Results

### Expression of neurotrophic factors in human bladder cancer suggests a critical role for nociceptors

To identify candidate tumor types whose pathogenesis may be influenced by nociceptors, we used the KM-plotter survival analysis tool^25^, which integrates gene expression and clinical outcome data from the GEO, EGA, and TCGA databases, to ask whether mRNA expression of nociceptor-associated genes correlates with a distinct prognosis in each of 22 major human malignancies. Since the low absolute amount of mRNA present within axons precludes direct detection of nociceptor-derived transcripts in tumor biopsies, we focused instead on neurotrophic factors that promote sensory innervation and sensitization, including NGF (nerve growth factor)^26^, BDNF (brain-derived neurotrophic factor)^27^, and GDNF (glial cell line-derived neurotrophic factor)^28^. Consistent with a recent report^23^, high expression of each of these cytokines correlated inversely with survival in gastric cancer (**Figure 1A**). The only other malignancy that showed a similar inverse correlation between survival and high expression of all three neurotrophins was bladder cancer (BC). Indeed, high expression of *NGF* was associated with a ∼four-fold decrease in median survival as compared to BC with low *NGF* expression (24 months vs. 106 months, respectively), suggesting that the ability of BC to promote sensory innervation may contribute to disease severity (**Figure 1B**).

**Figure 1.**
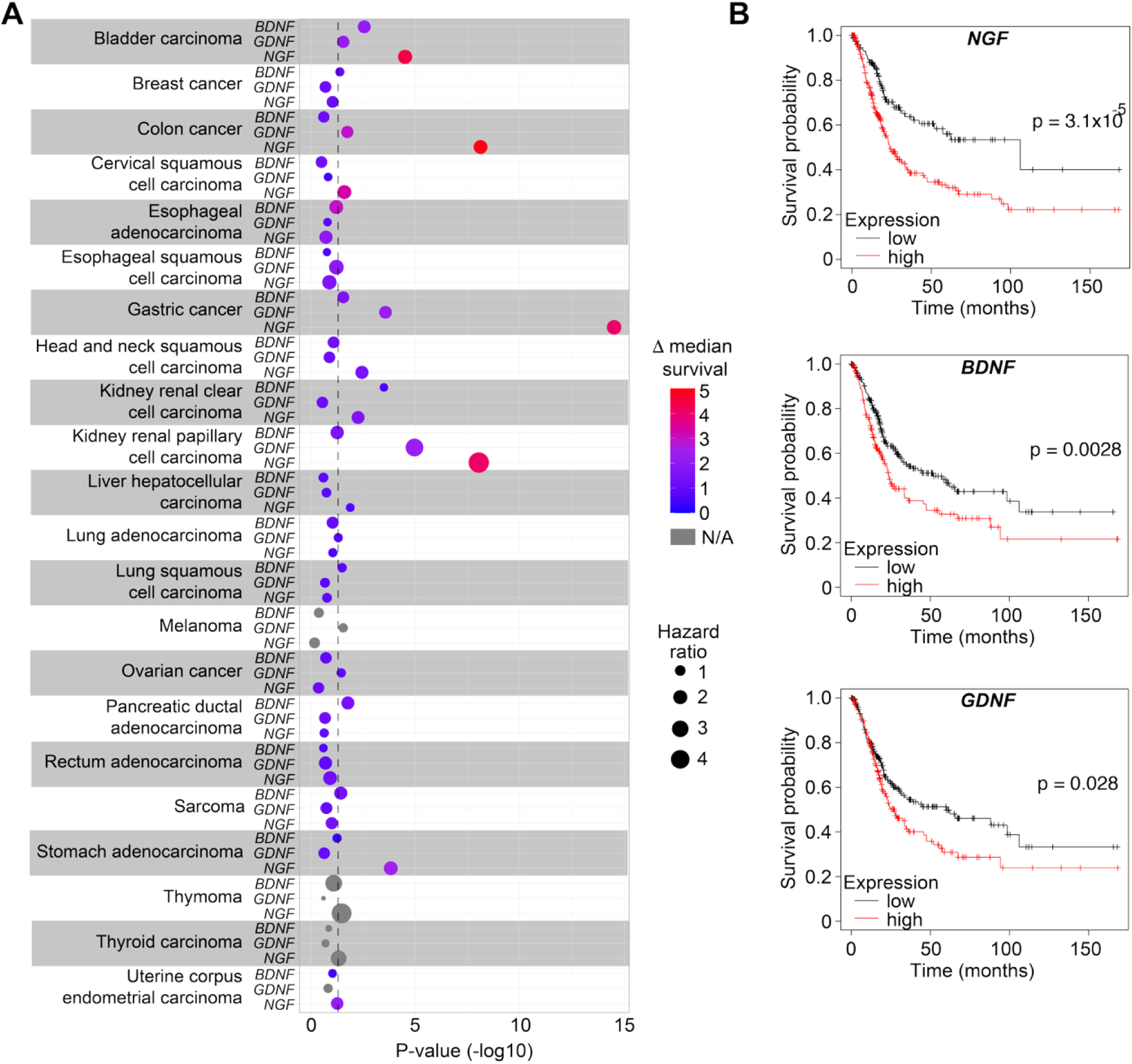
Expression of neurotrophins correlates with poor prognosis in bladder carcinoma patients. KMplotter was used to stratify patient populations with indicated malignancies based on the expression of *NGF, BDNF*, and *GDNF*. (**A**) Survival analysis was performed using the upper and lower quartiles (high and low expressors) for each gene, and the median survival difference was calculated for curves that dropped below 50%. (**B**) Kaplan-Meier plots summarizing the effect of expression of the indicated neurotrophic factors in bladder carcinoma patients. Log-Rank test was used for statistical comparison between curves.

### Orthotopic bladder cancer formation in mice requires monocyte-derived MDCSs

To directly assess the role of nociceptors in the pathogenesis of BC, we used an orthotopic model of non-muscle-invasive bladder carcinoma (NMIBC), whereby MB49 tumor cells were administered via a catheter into the bladder of mice that had been preconditioned by intravesical instillation of poly-*D*-lysine^29–31^. Fourteen days after MB49 cell instillation, the majority of mice (84%) had developed sub-mucosal tumors of varying sizes, which had penetrated the transitional urothelium and underlying basement membrane, but without invading the muscularis layer (**Figure 2A** and **S1A**). To search for markers that can discriminate MB49 tumors from surrounding non-malignant bladder tissue, we performed mRNA sequencing of MB49 cells (**Figure S1B**). Flow cytometry confirmed at the protein level that *Cd34*, *Cd44*, *Itgb1*, and *Lgals3* are among the highest-expressed transmembrane markers on MB49 cells (**Figure S1C)**. Among these, CD44 was chosen as an MB49 marker that allowed clear identification of tumors in tissue sections by immuno-fluorescence microscopy (**Figure 2B**).

**Figure 2.**
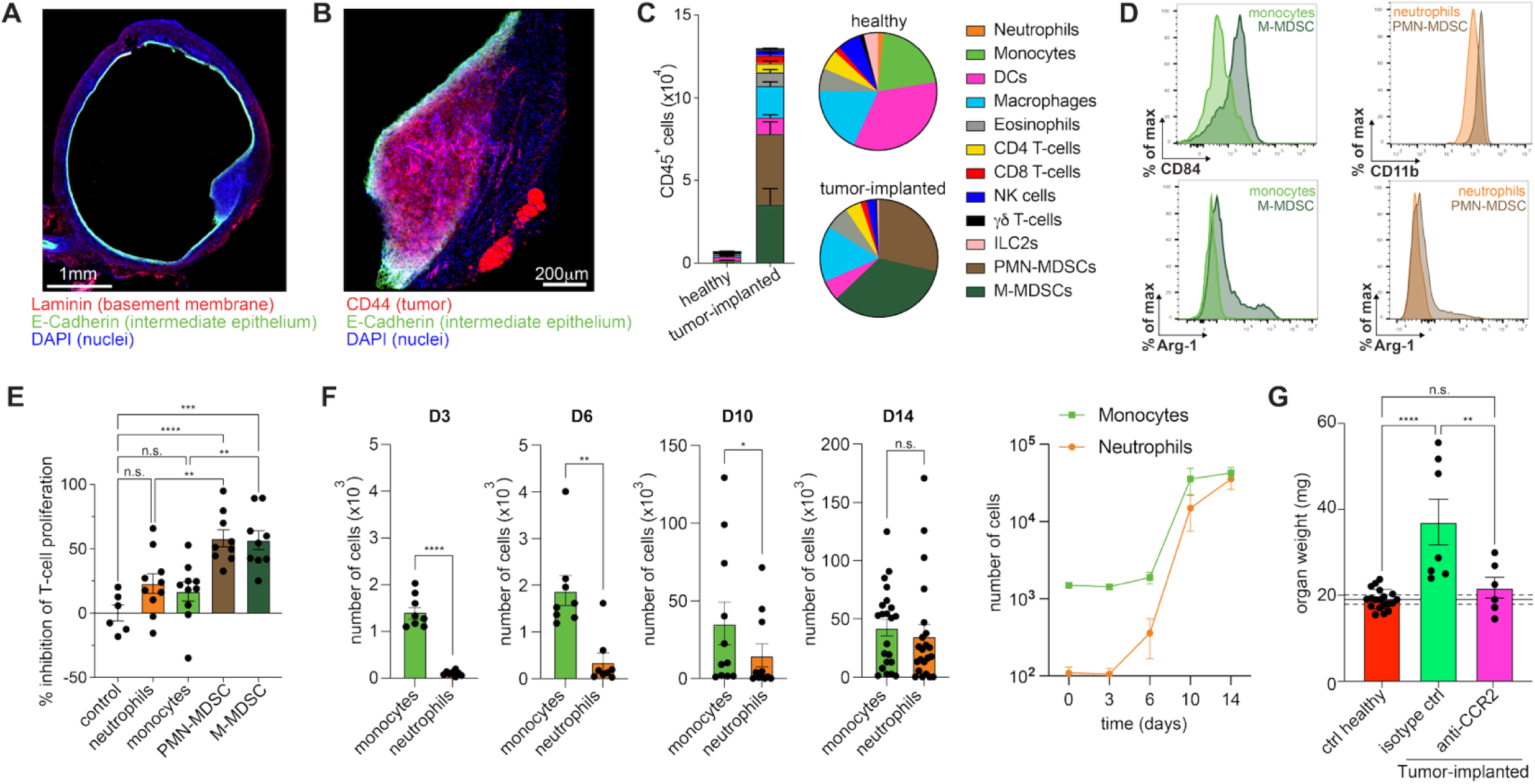
Early monocyte infiltration promotes tumor growth in orthotopic bladder cancer. (**A-B**) The bladder of a MB49 tumor-bearing mouse was harvested on day 14 after intravesical tumor cell injection, fixed, cryosectioned, stained with the indicated antibodies, and imaged on a confocal microscope. **(C-D)** Single-cell suspensions generated from bladders of healthy or MB49 tumor-bearing mice were analyzed by flow cytometry (day 14), and (**C**) indicated cell types were enumerated (n=22, error bars indicate SEM), or (**D**) the expression of indicated markers on monocytes and neutrophils from healthy bladders and tumor-derived M-MDSC and PMN-MDSC was assessed. (**E**) Myelopid cell subsets were purified from healthy spleens or tumor-bearing bladders and co-cultured with OVA-pulsed DCs and cell trace violet-loaded OVA-specific naive OT-I T-cells. T-cell proliferation was measured by assessing cell trace violet dilution by flow cytometry 3 days later. Means +/- SEM are shown, n=6-10. (**F**) Cells expressing phenotypic markers of monocytes and neutrophils were enumerated by flow cytometry in orthotopic MB49 tumors at the indicated time points. n=8-22 mice per group, error bars indicate SEM. (**G**) MB49-implanted mice were treated daily starting on day 4 with monocyte-depleting anti-CCR2 or isotype control antibody, and the weight of their bladders was measured on day 14 as a proxy for tumor growth. Means +/- SEM are shown, n=19 for healthy bladders and n=6-7 for tumor-implanted mice. Dashed lines indicate the 95% confidence interval for the weight of healthy bladders. One-way ANOVA with Tukey’s multiple comparisons test (**C,E**) or Welch’s t-test (**D**) were used for statistical analysis. *p<0.05, **p<0.01, ***p<0.001, ****p<0.0001.

Next, we set out to characterize the neuro-immune landscape in normal and tumor-bearing bladders. Compared to healthy bladders, tumor-bearing bladders harbored a large infiltrate of CD45^+^ immune cells (**Figure S1D**) that were further characterized by flow cytometry of single-cell suspensions (**Figure S1E**). The immune compartment of healthy bladders was dominated by classical dendritic cells (cDCs), monocytes, and macrophages, which, together, accounted for ∼75% of all CD45^+^ cells (**Figure 2C**). In day 14 tumor-bearing bladders, myeloid leukocytes were even more abundant (>80% of CD45^+^ cells), primarily due to a marked increase in CD11b^+^ cells with phenotypic characteristics of monocytes (Ly6C^+^ Ly6G^-^) or neutrophils (Ly6C^int^ Ly6G^+^).

Within the tumor microenvironment, leukocytes expressing the Ly6C or Ly6G markers are usually equated with monocytic (M-) or polymorphonuclear (PMN-) myeloid-derived suppressor cells (MDSCs), respectively^7^. While the unambiguous distinction of MDSCs from other myeloid cell types remains a challenge, the expression of known MDSC markers – CD84 on Ly6C^+^ cells^32^ and upregulation of CD11b on Ly6G^+^ cells^7^, as well as the expression of Arg-1 in both subsets^33^ (**Figure 2D**), suggested they represent *bona fide* MDSCs. Accordingly, these cells exhibited an altered cellular morphology. Specifically, M-MDSCs presented with a foamy appearance and were larger in size compared to splenic or healthy bladder monocytes, while PMN-MDSCs contained more hypersegmented nuclei, compared to bone marrow or splenic neutrophils (**Figure S1F**), as reported previously^34^. To formally test these putative MDSCs for their immunosuppressive ability, we co-cultured ovalbumin (OVA)-laden DCs with naïve, OVA-specific (OT-I) CD8^+^ T-cells in the presence or absence of FACS-purified, tumor-derived MDSCs or control splenic monocytes or neutrophils. Indeed, both tumor-derived myeloid cell subsets inhibited T-cell proliferation, confirming their immunosuppressive capacity and their likely identity as MDSCs **(Figure 2E)**.

To better understand the dynamics of TME development after BC seeding, we next isolated the bladders of mice at 3, 6, and 10 days post tumor implantation and analyzed their immune infiltrate. While neutrophils and neutrophil-derived cells were rare in healthy bladders and remained very sparse until day 10 after tumor challenge, monocytes and monocyte-derived cells were readily detectable in steady-state bladders and rapidly accumulated upon tumor implantation (**Figure 2F**). To assess the role of monocytes and their offspring in BC tumorigenesis, tumor-challenged mice were systemically treated with an anti-CCR2 MAb^35,36^, which depletes Ly6C^Hi^ monocytes, the putative precursors of M-MDSCs, for ∼5 days^37^. Bladders were harvested on day 14, and tumor burden was assessed by comparing their weight to tumor-free control bladders. Bladder weight was significantly increased in MB49-challenged monocyte sufficient mice, and these bladders contained macroscopically visible tumors. By contrast, when mice were treated with anti-CCR2 starting on day 4 after tumor challenge, bladders retained a normal weight and no tumors were detected, indicating that early monocyte recruitment plays an essential role in the establishment of MB49 bladder carcinoma (**Figure 2G**).

### Bladder cancer formation depends on tumor-induced nociceptor innervation

Having characterized the immune infiltrate of tumor-bearing bladders, we next set out to determine whether and to what extent bladder tumors receive nociceptive innervation. To unambiguously identify nociceptive fibers, we analyzed *Scna10^Cre/+^ R26^lsl-TdTomato^* mice (henceforth referred to as NaV1.8^TdT^ mice), which express TdTomato as a fluorescent reporter in NaV1.8^+^ nociceptors^38^. Immunofluorescence imaging of whole-mounts of healthy murine bladders identified a dense network of nociceptive fibers immediately underneath the uroepithelial barrier **(Figure S2A** and **Movie S1)**. Similarly, when bladders were harvested on day 14 after tumor implantation, subjected to tissue clarification using the iDISCO protocol^39^ and imaged by confocal microscopy, MB49 tumors were densely innervated by nociceptors (**Figure 3A** and **Movie S2**), suggesting that the growing tumors may induce axonal sprouting and neo-neurogenesis. Accordingly, tumor cells established close contacts with nociceptors when co-cultured *in vitro* (**Figure S2B**) and the presence of MB49 cells or MB49-conditioned medium promoted axonogenesis in freshly cultured nociceptors (**Figure 3B** and **S2C**). Furthermore, consistent with our analysis of human BC (**Fig. 1**), an RNAseq analysis showed that MB49 cells express *Ngf* as well as *Gdnf* and *Bdnf* (**Figure S2D**). Indeed, the ability of MB49 cells to promote axonogenesis was significantly reduced after deletion of the *Ngf* gene - a defect that could be rescued by lentiviral reconstitution of *Ngf* expression (**Figure 3B**). Moreover, in co-cultures, MB49 cells sensitized nociceptors, resulting in an increased release of the neuropeptide CGRP upon stimulation with capsaicin, a TRPV1 channel agonist, (**Figure S2E** and **3C**). Collectively, these data indicate that BC cells engage in active communication with nociceptors to promote axonal sprouting, at least in part, by releasing NGF and to modulate the neuronal response to activating stimuli.

**Figure 3.**
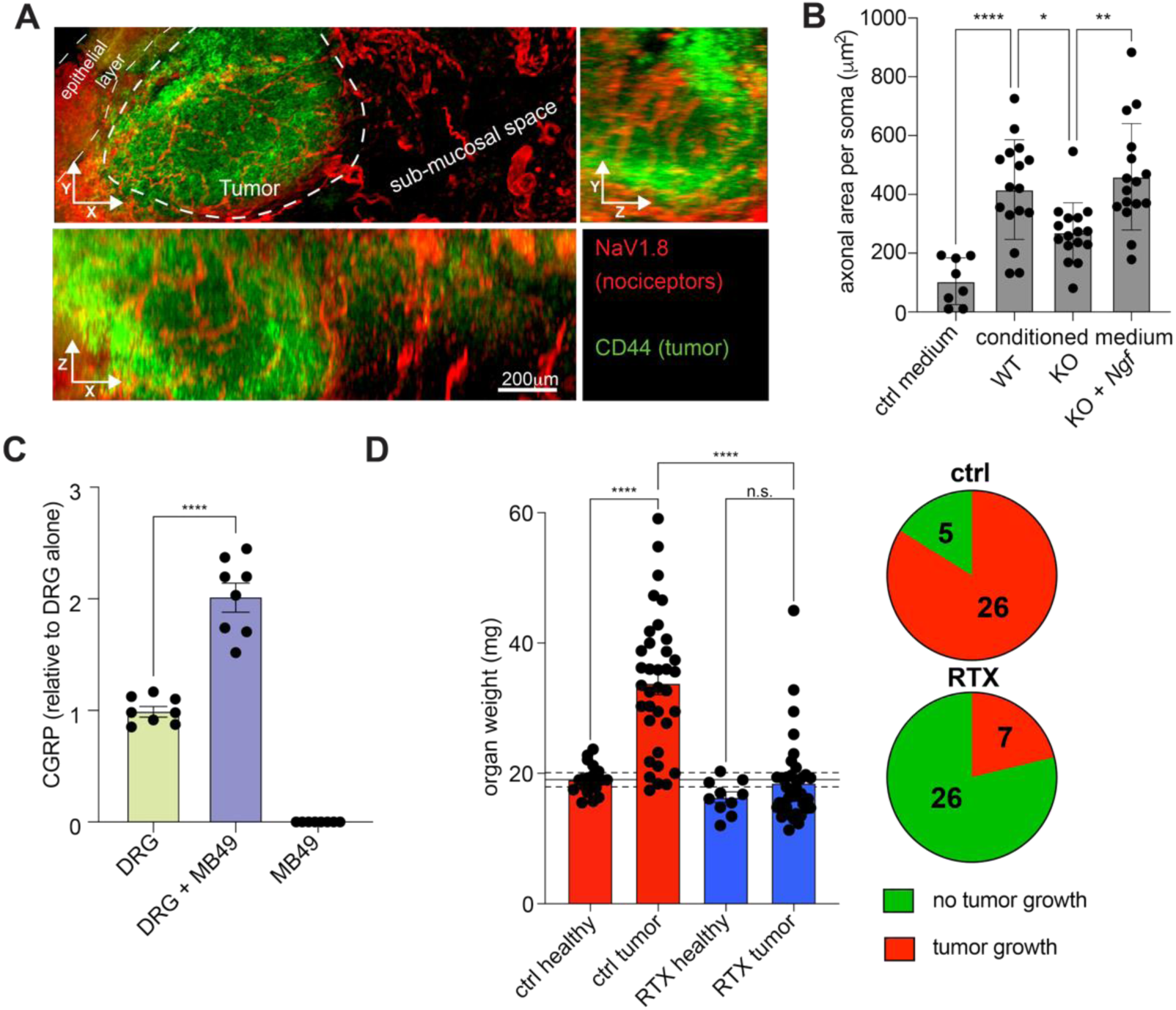
Nociceptive innervation is required for efficient formation of orthotopic bladder cancer. (**A**) MB49 tumors in bladders of NaV1.8^TdT^ mice were harvested on day 14 after implantation, iDISCO-clarified, stained with the indicated antibodies, and analyzed using confocal microscopy. (**B**) Freshly harvested, de-axonized nociceptors were prepared from DRGs, cultured for 16 hours alone or in the presence of medium conditioned by WT, *Ngf*-KO, or *Ngf*-reconstituted *Ngf-*KO MB49 cells, fixed, stained with β3 tubulin antibody, and analyzed by confocal microscopy. The total area of sprouted axons per soma was quantified using an automated script in Fiji. Mean +/- SEM of 4 independent experiments is shown. (**C**) 7 days after plating, nociceptors were co-cultured with MB49 cells or control medium for 24 hours, and treated with capsaicin for 2 hours. CGRP release was measured by ELISA. Mean +/- SEM of 3 independent experiments is shown. (**D**) Bladders of nociceptor-sufficient (ctrl) and –depleted (RTX) mice were implanted with MB49 cells, and their weight was measured on day 14 as a proxy for tumor growth. Mean +/- SEM of 10-19 healthy and 31-33 tumor-bearing mice per group are shown. Dashed lines indicate the 95% confidence interval for the average weight of healthy bladders. One-way ANOVA with Tukey’s multiple comparisons test (**B**), Welch’s t-test (**C**), or two-way ANOVA with Tukey’s multiple comparisons test (**D**) were used for statistical analyses. *p<0.05, **p<0.01, ***p<0.001, ****p<0.0001.

To formally test the pathophysiological relevance of these *in vitro* observations and to assess the impact of nociceptors in BC tumorigenesis, we used resiniferatoxin (RTX)^40^, a TRPV1 superagonist, to selectively ablate TRPV1^+^ nociceptors in mice prior to MB49 cell instillation. RTX treatment did not detectably alter the uroepithelial lining (**Figure S3A**), overall morphology (**Figure S3B**), or the immune landscape in steady state bladders (**Figure S3C**). However, nociceptor-depleted mice developed tumors at dramatically lower rates. While 84% (26/31) of MB49-implanted nociceptor-competent mice developed tumors within 2 weeks, the tumor incidence in nociceptor-ablated mice was only 21% (7/33) (**Fig. 3D**).

### Nociceptors prevent bladder cancer rejection by cytotoxic T cells

Tumor rejection is usually mediated by CD8^+^ CTLs and/or NK cells^41^. While NK cells can kill tumor cells in an antigen-independent manner, activation of a CTL response requires the presentation of tumor-specific antigens to naïve CD8^+^ T-cells within tumor-draining lymph nodes. Indeed, the cellularity of bladder-draining lymph nodes of tumor-challenged mice was significantly increased in both the nociceptor-competent and -ablated animals, presumably reflecting a recent or ongoing immune response against tumor antigens (**Figure 4A**). Accordingly, using the expression of the inducible co-stimulator (ICOS) as a marker of T-cell activation^42^, we observed that tumor challenge in both nociceptor-ablated and control mice resulted in a similar increase in the proportion of activated CD8^+^ T-cells (**Figure 4B**) as well as, to a lesser extent, CD4^+^ T-cells (**Figure S4**).

**Figure 4.**
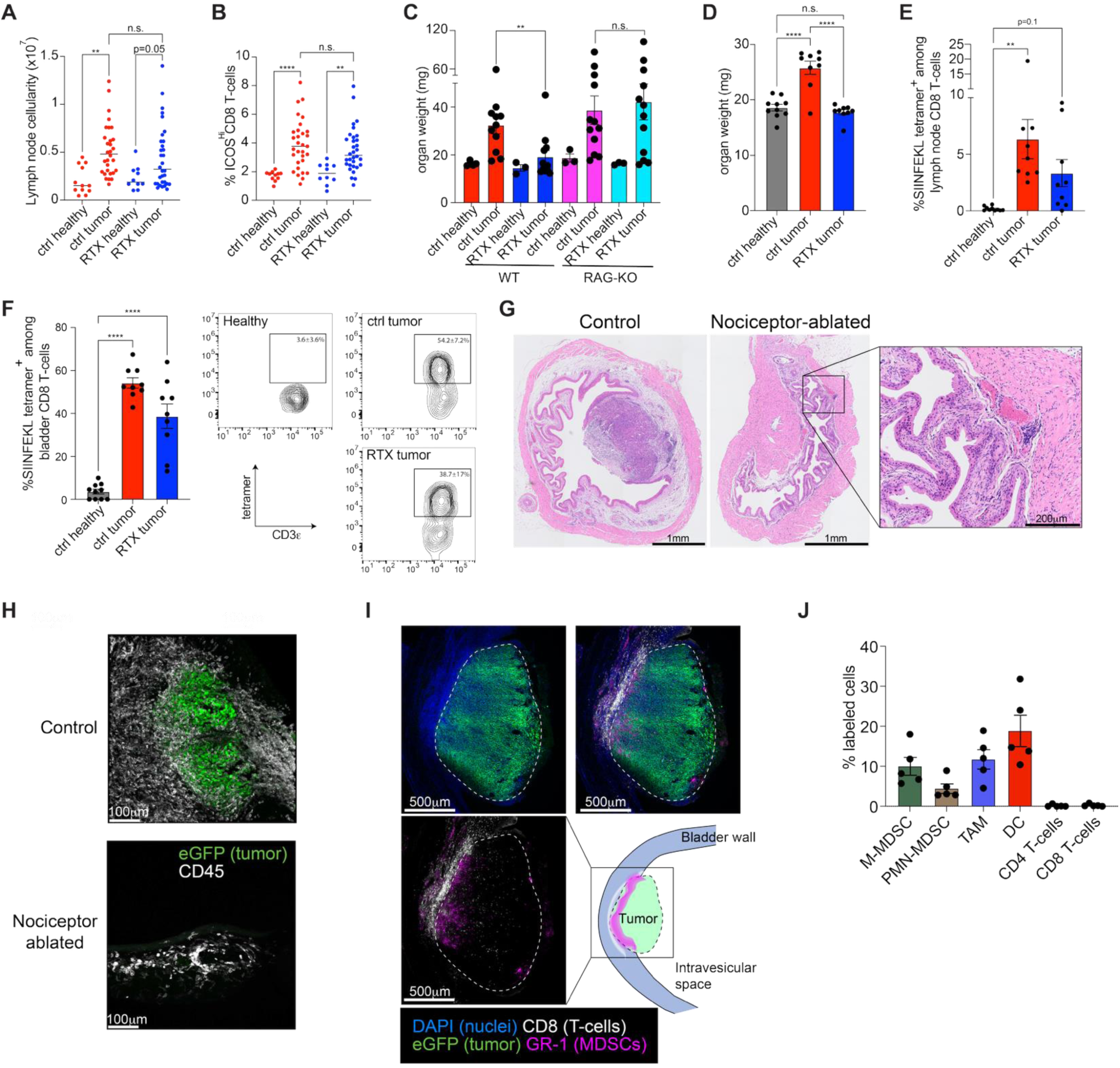
Nociceptors enable bladder cancer formation by preventing CTL-mediated tumor rejection. (**A,B**) The bladder draining lymph nodes of healthy or day 14 tumor-implanted nociceptor-ablated (RTX) and -competent (ctrl) mice were harvested and analyzed by flow cytometry to assess (**A**) total cellularity and (**B**) the frequency of ICOS^+^ CD8 T-cells. The data represent 10-12 healthy and 30-37 tumor-implanted animals per group. (**C**) The weight of bladders of healthy or tumor-implanted nociceptor-ablated (RTX) and -competent (ctrl) WT and RAG2-deficient mice was measured on day 14 as a proxy for tumor growth. Mean +/- SEM of 3 healthy and 11-12 tumor-bearing mice per group is shown. (**D-F)** Nociceptor-competent (ctrl) and –ablated (RTX) mice were implanted with SIINFEKL-expressing MB49 cells, and the organ weight (**D**), proportion of SIINFEKL-specific CD8 T-cells within the lymph nodes (**E**) and bladders (**F**) was assessed on day 14. Representative examples of FACS plots and mean +/-SEM of 9-10 animals per group are shown. (**G**) Bladders of tumor-implanted nociceptor-ablated (RTX) and -competent (ctrl) mice were analyzed after H&E staining. The inset shows scar tissue bordered by a cluster of immune cells. (**H**) eGFP-MB49 cells were implanted in nociceptor-ablated (RTX) and -competent (ctrl) mice, and the bladders were harvested on day 14, fixed, cryo-sectioned, stained with CD45 antibodies, and analyzed by confocal microscopy. (**I**) Bladders of nociceptor-competent eGFP-MB49-implanted mice were harvested on day 14, fixed, cryo-sectioned, stained with the indicated antibodies, and analyzed by confocal microscopy. (**J**) Bladders of sLP-mCherry-MB49-implanted mice were harvested on day 14, and the acquisition of sLP-mCherry in the indicated immune cell subsets was assessed by flow cytometry. Data represent mean +/- SEM of cells from 5 tumor-bearing animals. For statistical analysis, a two-way ANOVA with Tukey’s multiple comparisons test (**A,B**), Welch’s t-test (**C**), or one-way ANOVA with Tukey’s multiple comparisons test (**D-F**) were used. *p<0.05, **p<0.01, ***p<0.001, ****p<0.0001.

Importantly, the protective effect of nociceptor ablation was completely lost in *Rag2*-deficient (Rag-KO)^43^ mice, which are devoid of B and T-cells (**Figure 4C**), indicating that the prevention of tumor growth in nociceptor-ablated WT animals was a result of an improved adaptive anti-tumor immune response, presumably by CTL, rather than altered urothelial barrier function or other non-immune effects. Thus, to dissect the antigen-specific T-cell response in MB49 tumors, we generated MB49 cells expressing SIINFEKL, the immunodominant epitope of chicken ovalbumin (OVA) for CD8 T cells in C57BL6 mice^44,45^. Like their WT counterparts, SIINFEKL-MB49 cells readily gave rise to tumors in most nociceptor-competent mice but not in RTX-treated mice (**Figure 4D**). Notwithstanding, intravesical SIINFEKL-MB49 cell challenge induced the accumulation of SIINFEKL-specific T-cells in both bladder-draining lymph nodes (**Figure 4E**) and bladders (**Figure 4F**) of nociceptor-competent and nociceptor-ablated animals. SIINFEKL-specific T cells were virtually absent from the bladders of tumor-naïve control mice. Taken together, these data demonstrate that MB49 bladder tumors induce an adaptive, tumor antigen-specific CTL response in bladder-draining lymph nodes irrespective of whether nociceptors are present or not, but this T cell response appears to be only capable of eliminating bladder tumors after nociceptor ablation.

Although nociceptors innervate peripheral lymph nodes^38^ and regulate antigen transport to lymph nodes by DCs^9^, the fact that tumor-induced T cell responses occurred similarly in the presence and absence of nociceptors implied that the pro-tumorigenic effect of nociceptors likely occurred within the bladder, rather than in draining lymph nodes. Indeed, hematoxylin and eosin (H&E) staining of bladders on day 14 after tumor implantation revealed that tumors in control animals exhibited the characteristic morphology of urothelial adenocarcinoma, accompanied by pronounced tissue edema, but without muscularis invasion (**Figure 4G**). In contrast, although most bladders of nociceptor-ablated animals showed no obvious signs of tumor growth, these tumor-free organs, unlike healthy control bladders, contained subepithelial foci of fibrotic tissue that were frequently bordered by aggregates of mononuclear leukocytes.

These scar-like lesions presumably represented the sites where MB49 cells had established an initial foothold before being eradicated by the ensuing CTL response. To better differentiate tumors vs. healthy tissue and to ascertain whether nociceptor-ablated bladders are truly devoid of all tumor cells, we implanted MB49 cells expressing enhanced green fluorescent protein (eGFP) into control and nociceptor-ablated hosts. On day 14, eGFP^+^ lesions were readily detectable in control animals, while eGFP^+^ cells were entirely absent from most nociceptor-ablated bladders (**Figure 4H**). The latter contained CD45^+^ immune cell aggregates resembling those that were observed in H&E stained sections adjacent to scar tissue. Thus, by day 14, tumor cells had been completely eradicated, presumably by CTL. By contrast, in nociceptor-sufficient bladders, tumors exhibited an immune-excluded phenotype^46^; the vast majority of CD45^+^ leukocytes infiltrating the tumor mass were GR1^+^ myeloid cells, which were particularly abundant at the tumor interface with the bladder wall (**Figure 4I**). There was also a substantial population of CD8 T cells within the bladder wall; however, CD8^+^ CTLs were almost exclusively positioned at the tumor margin and appeared to be excluded from the tumor itself. The exclusion of CD8^+^ T-cells from the TME was further confirmed by the use of a tumor niche-labeling approach. This strategy relied upon transfected MB49 tumor cell secretion of liposoluble-tagged mCherry (sLP-mCherry) which accumulates in nearby tumor-associated cells^47^. Consistent with our histological findings, the sLP-mCherry signal was primarily detected in M-MDSCs, TAMs, and DCs, but not in T-cells (**Figure 4J**).

In aggregate, the above results demonstrate that MB49 tumors harbor an immune infiltrate dominated by MDSC as well as dense nociceptive innervation, both of which are indispensable for optimal tumor establishment. These observations suggest that a three-way cross-talk between tumor, nociceptors, and monocytes may drive the establishment of an immunosuppressive TME, which protects the tumor against CTL-mediated eradication.

### Nociceptors promote monocyte differentiation into M-MDSC

Monocytes are short-lived, circulating myeloid leukocytes comprising classical (inflammatory) and non-classical (patrolling) subsets based on their differential expression of Ly6C and other markers^48^. Ly6C^Lo^ non-classical monocytes mainly act within the vasculature, while the Ly6C^Hi^ classical monocytes are recruited to peripheral tissues, where they can assume phenotypic and functional features of macrophages or DCs^48^. Blood-borne monocytes are unlikely to interact with nociceptors whose projections are not thought to reach into the intravascular space, whereas classical monocytes that extravasate into densely innervated bladder tumors would likely be exposed to nociceptors. Although nociceptors are known to regulate the function of other myeloid leukocytes, including DCs and macrophages^9,10,15,16^, the mechanisms and consequences of nociceptor interactions with monocytes are not well understood.

To address this question, we compared the transcriptomes of purified splenic Ly6C^Hi^ monocytes that were either cultured alone or cocultured with primary murine nociceptors in the presence or absence of LPS stimulation (**Figure 5A**). In agreement with previous studies describing anti-inflammatory effects of nociceptors on macrophages^49^, the presence of nociceptors dampened the LPS-induced expression of inflammatory cytokines, including TNFα, IL-6, and IL-12, while upregulating the anti-inflammatory cytokine IL-10 (**Figure S5A**). Notably, the mere presence of nociceptors, without any further stimulus, was sufficient to reprogram the transcriptome of monocytes (**Figure 5A,B**), a finding similar to our previous observations in DCs^9^. Gene ontology (GO) term enrichment analysis^50,51^ revealed that nociceptors upregulated in monocytes pathways involved in cell migration and motility, while immune and defense response pathways were downregulated (**Figure S5B**), indicating that nociceptors induce an anti-inflammatory monocyte program. Indeed, the highest upregulated gene in monocytes cocultured with nociceptors was *Arg1* (**Figure 5B**), a gene encoding the immunosuppressive enzyme arginase 1 (Arg-1). Among leukocytes, Arg-1 is mainly expressed by alternatively activated (M2) macrophages, tolerogenic DCs, and MDSCs^52^.

**Figure 5.**
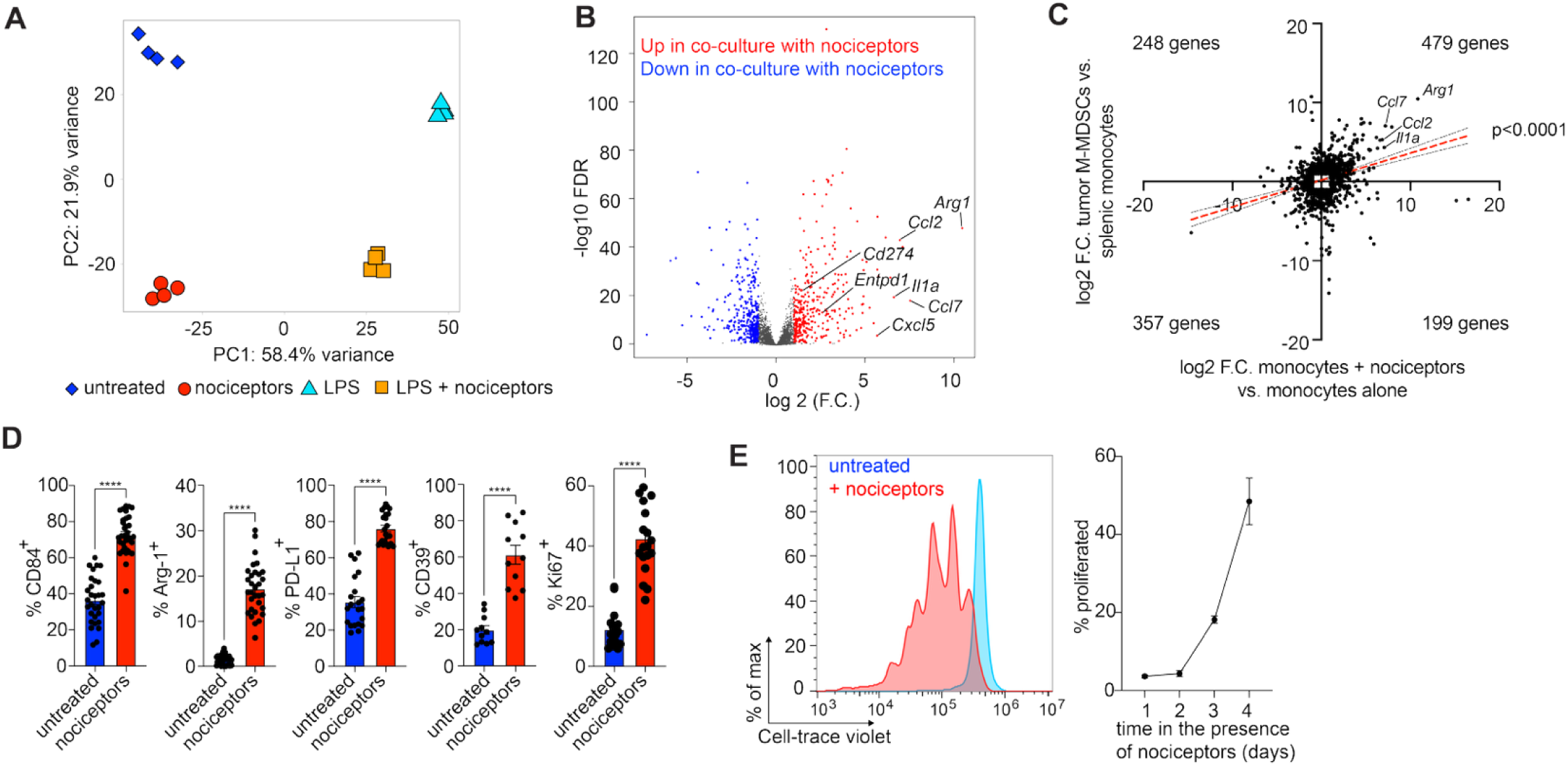
Nociceptors induce an MDSC-like phenotype in cocultured monocytes. (**A**) FACS-purified Ly6C^Hi^ splenic monocytes were cultured overnight alone or with nociceptors in the presence or absence of LPS, and analyzed by RNAseq. Principal component analysis (**A**) and volcano plot comparing untreated monocytes cultured alone or with nociceptors (**B**) are shown. Selected genes associated with MDSC functions are highlighted. (**C**) FACS-purified MB49 tumor-derived M-MDSCs and splenic monocytes were analyzed by RNAseq. Genes that were differentially regulated between M-MDSCs and monocytes (X-axis) or monocytes co-cultured with nociceptors and monocytes cultured alone (Y-axis) were plotted, and linear regression was performed to assess the correlation between the datasets. (**D**) FACS-purified splenic Ly6C^Hi^ monocytes were co-cultured overnight with nociceptors, and the expression of indicated markers was assessed by flow cytometry. Mean +/- SEM of 5-13 independent experiments is shown. Welch’s t-test was used for statistical analysis. (**E**) FACS-purified splenic Ly6C^Hi^ monocytes were labeled with Cell-trace violet and co-cultured with nociceptors for the indicated period. Divided cells were identified based on Cell-trace violet dilution and enumerated. Mean +/- SEM of 2 independent experiments (right), and a representative histogram of a 4-day culture (left) are shown. *p<0.05, **p<0.01, ***p<0.001, ****p<0.0001.

Monocytes can give rise to a variety of functionally distinct progeny, including macrophages, DC-like cells, and M-MDSCs; however, the pathways that drive M-MDSC differentiation are incompletely understood, and the distinction between MDSCs and other myeloid cell types is often ambiguous^53^. Nevertheless, monocytes cocultured with nociceptors did not upregulate common macrophage markers, e.g. *Adgre1* (encoding F4/80) or *Cd68*, nor did they express genes involved in the differentiation and functions of DCs (*Zbtb46, Itgax, H2-Aa, H2-Ab1, Cd1d1, Cd1d2, Cd80, Cd83*) or M2 macrophages (*Cebpb*, *Pparg*), even though some M2 associated markers (*Cd163*, *Msr1* (encoding CD204), *Mrc1* (encoding CD206)) were upregulated, as was *Trem2*, which is reportedly expressed by both M-MDSCs^54^, and a subset of monocyte-derived macrophages^55^ (**Figure S5C**).

By contrast, nociceptor exposure induced in monocytes multiple transcripts that have been implicated in MDSC functions^56^, including *Il1a*^57^*, Ccl2*^58^*, Ccl7*^59^*, Cxcl5*^60^*, Entpd1*^61^*, and Cd274*^62^ (**Figure 5B** and **S5C**). Indeed, the transcriptional profile of monocytes co-cultured with nociceptors was reminiscent of *bona fide* M-MDSCs isolated from MB49 tumors (**Figure 5C**), and gene-set enrichment analysis (GSEA)^63,64^ confirmed that nociceptor-conditioned monocytes were transcriptionally more similar to tumor-derived M-MDSCs than to splenic monocytes.

Conversely, tumor-derived M-MDSCs were more akin to monocytes co-cultured with nociceptors compared to monocytes cultured alone (**Figure S5D**). These transcriptional changes were further corroborated at the protein level; a flow cytometry analysis showed upregulation of Arg-1, PD-L1 (encoded by *Cd274*), and CD39 (encoded by *Entpd1*), as well as CD84, the only definitive M-MDSC marker identified to date^32^ (**Figure 5D**). Of note, although monocytes are considered post-mitotic and short-lived^65^, monocytes exposed to nociceptors upregulated the proliferation marker Ki-67 (**Figure 5D**) and proliferated vigorously (**Figure 5E**).

This monocyte response was a unique effect of their encounter with nociceptors since monocyte co-culture with a stromal cell line, OP9^66^, did not induce appreciable phenotypic changes (**Figure S6A**). Moreover, when nociceptors were killed by fixation, they lost their ability to reprogram monocytes (**Figure 6A**). Additionally, MDSC differentiation was only observed with classical monocytes. When Ly6C^Lo^ non-classical monocytes were cocultured with nociceptors, they upregulated PD-L1 but not Arg-1, and they expressed little or no detectable CD84 (**Figure S6B**). Thus, nociceptor-mediated reprogramming of Ly6C^Hi^ monocytes toward an M-MDSC phenotype is the result of an active and subset-specific dialogue between these two cell types.

**Figure 6.**
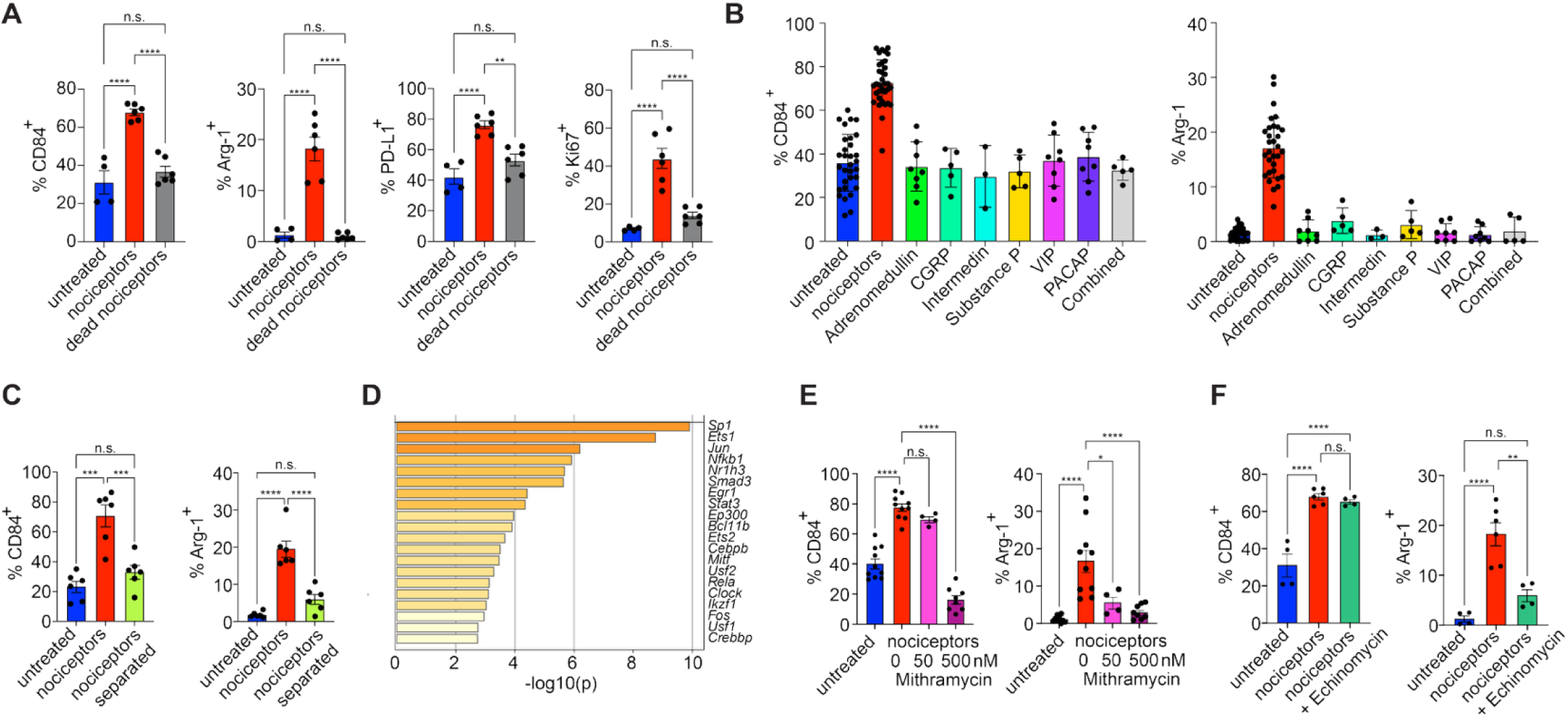
Nociceptors drive monocyte differentiation by activating the Sp1 and Hif1α pathways. FACS-purified splenic Ly6C^Hi^ monocytes were (**A**) cultured alone or co-cultured overnight with live or PFA-fixed nociceptors, or (**B**) treated with the indicated neuropeptides (1μM). (**C**) Monocytes and nociceptors were cultured in a Transwell plate either in the same compartment or in different compartments separated by a filter membrane to allow or prevent physical contact, respectively. **(D)** TRRUST analysis of RNAseq data comparing FACS-purified Ly6C^Hi^ splenic monocytes cultured in the presence or absence of nociceptors. (**E-F**) FACS-purified splenic Ly6C^Hi^ monocytes were co-cultured with nociceptors in the presence or absence of (**E**) mithramycin (50 and 500nM) or (**F**) echinomycin (5nM), and the expression of indicated markers was assessed by flow cytometry. Mean +/- SEM of 2-4 independent experiments is shown. One-way ANOVA with Tukey’s multiple comparisons test was used for statistical analysis. *p<0.05, **p<0.01, ***p<0.001, ****p<0.0001.

### Nociceptor-induced M-MDSC differentiation is independent of neuropeptides

Neuropeptides are often considered the main modality by which nociceptors communicate with immune cells^67^. CGRP, in particular, has a potent immunosuppressive effect on macrophages^15^, neutrophils^13^, T-cells^19^, and innate lymphoid cells^68^. Additionally, recent work has shown that CGRP induces thrombospondin-1 (TSP1) in macrophages and neutrophils, a cytokine that promotes tissue repair^16^. Accordingly, an RNAseq analysis of CGRP-treated monocytes revealed that *Thbs1* (which encodes TSP1) was among the highest upregulated genes; however, CGRP treatment did not recapitulate the monocyte response to nociceptors and resulted in a comparatively smaller number of differentially regulated genes (**Figure S6C**).

While these findings indicated that CGRC by itself is insufficient to promote monocyte–MDSC differentiation, they did not rule out that CGRP contributes to BC formation by other means. Indeed, in a mouse model of subcutaneous (SQ) melanoma, nociceptor-derived CGRP was shown to promote tumor growth by inducing an exhausted phenotype in tumor-infiltrating CTL^19^. Yet another tumor-promoting activity of CGRP was identified in gastric cancer where this neuropeptide acts directly on tumor cells^23^. Thus, to rigorously explore the role of CGRP in BC, we implanted MB49 tumors in the bladders of CGRPα-deficient (*Calca^eGFP/eGFP^*) mice^69^. When compared to WT hosts, there was no difference in tumor growth or in the accumulation of immunosuppressive myeloid cells (**Figure S6D**), indicating that the tumor-promoting activity of nociceptors in BC involves mechanisms that may be distinct from other tumor types and/or anatomic locations.

To explore the monocyte response to nociceptive neuropeptides more broadly, monocytes were exposed to a panel of common agonists, including CGRP, adrenomedullin, intermedin, substance P, vasoactive intestinal peptide (VIP), and pituitary adenylate cyclase-activating polypeptide (PACAP). None of these agents alone or in combination induced CD84 or Arg-1 expression (**Figure 6B**).

Most neuropeptides signal through G protein-coupled receptors (GPCRs) to activate adenylyl cyclase (AC), resulting in the intracellular accumulation of cyclic adenosine monophosphate (cAMP). This effect can be mimicked pharmacologically by activating AC with forskolin or by exposure to 8-Br-cAMP, a synthetic cAMP analog^9^. However, neither of these compounds induced CD84 or Arg-1 upregulation in monocytes (**Figure S6E**), indicating that neuropeptides or other agonists of AC-linked GPCRs are insufficient to induce the M-MDSC phenotype.

This interpretation was also consistent with coculture experiments where monocytes and nociceptors were plated in different compartments of a Transwell device in which both cell types shared the same media, but physical interactions were prevented by a cell-impermeable filter membrane. In these culture conditions, monocytes failed to assume an MDSC phenotype (**Figure 6C**), suggesting that M-MDSC differentiation requires physical contact between monocytes and nociceptors or potentially involves secreted mediators that act only over short distances.

### Nociceptors drive M-MDSC differentiation of monocytes by activating the Sp1 and Hif1α pathways

Having ruled out neuropeptides as the primary drivers of monocyte–MDSC differentiation, we performed TRRUST (Transcriptional Regulatory Relationships Unraveled by Sentence-based Text mining)^70^ analysis of our RNASeq data to identify intracellular pathways underlying differential gene expression in monocytes alone *versus* monocytes cocultured with nociceptors. Comparing transcription factor signatures in the dataset, the analysis identified the Specificity protein 1 (Sp1) as the strongest candidate regulator of nociceptor-induced M-MDSC differentiation (**Figure 6D**). Accordingly, monocyte treatment with mithramycin, an Sp1 inhibitor, prevented the nociceptor-induced MDSC phenotype acquisition in a dose-dependent manner (**Figure 6E**). Of note, recent work has shown that Sp1 interacts with hypoxia-inducible factor 1α (Hif1α), whereby these two transcription factors promote each other’s effects on cancer-associated gene expression^71^. Indeed, upregulation of Arg-1, one of the key nociceptor-induced MDSC-associated genes, has previously been shown to be dependent on Hif1α^72,73^. Under normoxic conditions, Hif1α resides in the cytosol; it translocates to the nucleus to regulate gene expression during hypoxia^74^. Accordingly, in monocytes cultured under normoxic conditions in isolation, Hif1α was excluded from the nucleus. However, this nuclear exclusion of Hif1α was lost upon co-culture with nociceptors (**Figure S6E**), suggesting that the latter activate Hif1α in monocytes in a hypoxia-independent manner. Indeed, the Hif1α inhibitor echinomycin prevented nociceptor-induced Arg-1 upregulation in monocytes, without affecting CD84 expression (**Figure 6F**). Similarly, treatment of monocytes with a Hif1α activator, DMOG (dimethyloxalylglycine)^75^, induced Arg-1, but not CD84 upregulation (**Figure S6F**).

### The effect of nociceptors in MB49 tumors depends on anatomic context

The above observations are consistent with a model where nociceptors within developing bladder tumors promote the conversion of rapidly infiltrating monocytes into M-MDSCs, which subsequently protect the tumor against T-cell-mediated eradication. As discussed above, this tumor-promoting immunomodulatory activity of nociceptors in the bladder appears to be mechanistically distinct from that in SQ-implanted melanomas^19^ in terms of both target cell type (monocytes vs. CTL, respectively) and molecular pathway (contact-dependent Sp1/Hif1α agonism vs. secreted CGRP, respectively). To ask whether this distinction reflects differences in tumor type or anatomic environment, we assessed the impact of nociceptors on SQ-implanted MB49 tumors. Strikingly, nociceptor ablation had no discernible effect on the incidence or growth of SQ MB49 tumors, regardless of the number of implanted tumor cells (**Figure S7A**). A histologic examination of SQ MB49 tumors revealed the presence of nociceptors in peritumoral tissue, but not within the tumor itself, indicating that the TME in the SQ model does not faithfully recapitulate the composition of BC in its tissue of origin (**Figure S7B**).

In this context, it is pertinent that tumor-associated neutrophils can give rise to PMN-MDSC, which may also contribute to an immunosuppressive TME^76^. By day 14, when tumors were fully established, M-MDSC and PMN-MDSC had accumulated to similar frequencies in MB49 tumors in both the bladder (**Figure 2A**) and the subcutis (**Figure S7C)**. However, prior to day 10, M-MDSC far outnumbered PMN-MDSC in orthotopic bladder tumors (**Figure 2D**), whereas SQ tumors harbored similar numbers of the two MDSC subsets *a priori* (**Figure S7D**). Thus, we speculated that the immunosuppressive effect of neutrophils in the TME may be nociceptor-independent and, as a result, the rapid neutrophil recruitment into SQ lesions may obviate the need for nociceptor dependent M-MDSC formation. In support of this idea, an RNAseq analysis revealed that primary bone marrow neutrophils did not acquire an immunosuppressive phenotype when cocultured with nociceptors (**Figure S7E**). In fact, GO term enrichment analysis identified only relatively small nociceptor induced transcriptional changes in neutrophils, mostly in pathways involving lipid metabolism (**Figure S7F**). These findings suggest that in SQ tumors PMN-MDSC can arise rapidly in a nociceptor-independent fashion. By contrast, incipient tumors in the bladder do not initially recruit neutrophils, so nociceptor-mediated conversion of monocytes to M-MDSCs is critical for BC growth and persistence. This concept predicts that the pro-tumorigenic effect of nociceptors in BC may be limited to the earliest stages of tumor growth, prior to neutrophil entry.

### Rapid and selective monocyte recruitment to incipient bladder cancer lesions

To better understand the spatial relationships between tumor cells, nociceptors, and incoming monocytes during the first days after tumor seeding of the bladder, we performed confocal imaging of bladder whole mounts after intravesical instillation of eGFP-MB49 cells. By day 2 post-instillation, MB49 cells had penetrated across the urothelial layer and already initiated multiple contacts with bladder-innervating neurons (**Figure 7A** and **Video S3**). By day 4, the incipient tumor lesions had expanded and begun to attract numerous GR-1^+^ myeloid cells, which were dispersed throughout the tumor and frequently localized in immediate proximity to neuronal axons (**Figures 7A, S7G,** and **Video S4**). Importantly, the majority of the GR1^+^ cells also showed bright Ly6C staining, identifying them as monocytes, while Ly6C^Dim^ neutrophils were rarely detected (**Figure S7H).** By day 7, the tumor had assumed a rounded, nodular morphology with closely packed tumor cells and a dense GR1^+^ myeloid infiltrate. At this time point, isolated CD8^+^ CTL also began to appear; however, these T-cells were often found in the tumor periphery, and those that had accessed the TME were vastly outnumbered by GR-1^+^ cells (**Figure 7B** and **Video S5**).

**Figure 7.**
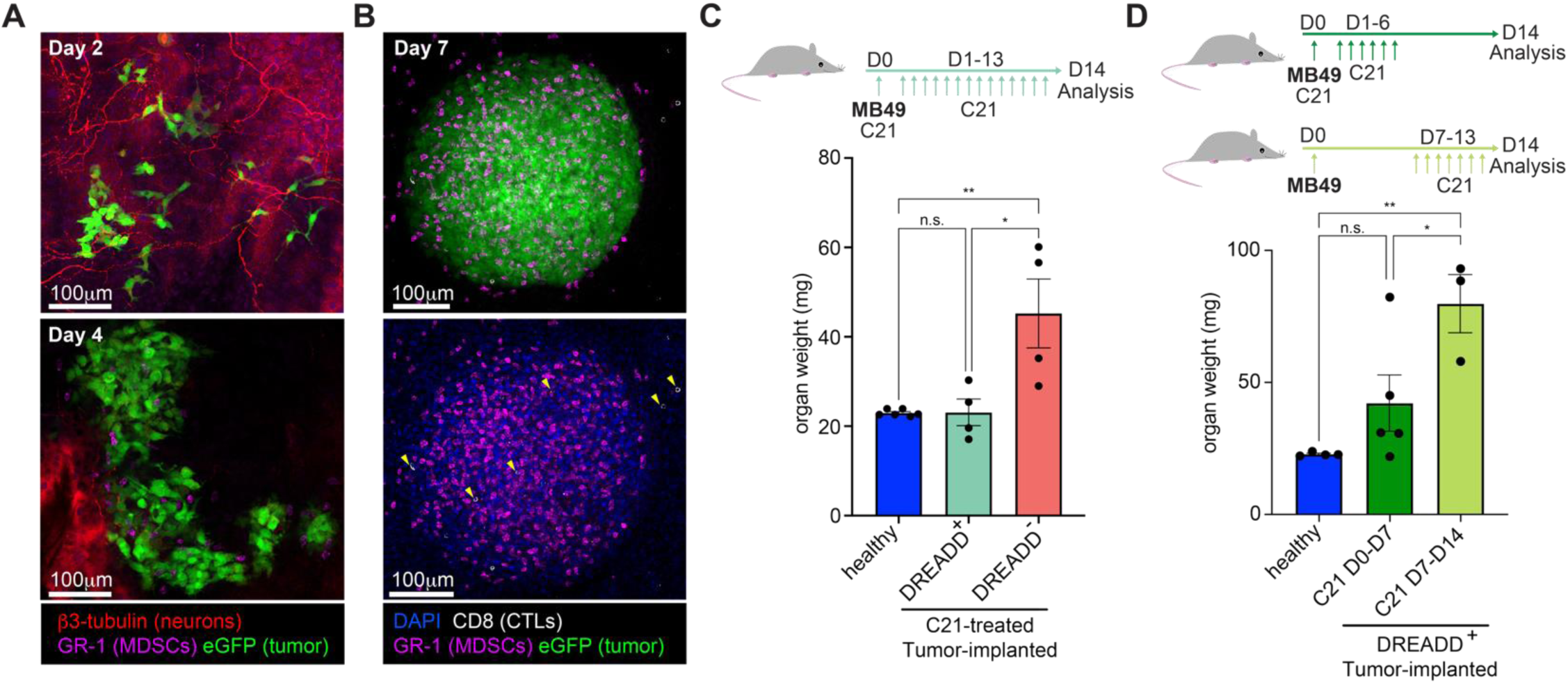
Nociceptor electrical activity is necessary during the early stages of MB49 tumor establishment. (**A,B**) Bladders of eGFP-MB49 tumor cell-implanted mice were harvested at the indicated time points, stained with the indicated antibodies, and analyzed as whole-mount preparations by confocal microscopy. CD8 T-cells are indicated with yellow arrows. (**C**) Bladders of NaV1.8*^Gi-DREADD^* mice or DREADD-negative littermate controls were implanted with MB49 tumors, and the mice received a daily injection of the DREADD agonist C21 (3mg/kg). Bladder weight as a proxy for tumor growth was assessed on day 14. A summary of 3 independent experiments is shown, n=4-6, error bars indicate SEM. (**D**) Bladders of NaV1.8*^Gi-DREADD^* mice were implanted with MB49 cells, and the animals received a daily injection of the DREADD agonist C21 during the early (D0-D6) or late (D7-D14) stage of tumor growth. Bladder weight as a proxy for tumor growth was assessed on day 14. A summary of 2 independent experiments is shown, n=3-5, error bars indicate SEM. One-way ANOVA with Tukey’s multiple comparisons test was used for all statistical analyses. *p<0.05, **p<0.01, ***p<0.001, ****p<0.0001.

### Nociceptors’ electrical activity is required for monocyte – MDSC differentiation in early bladder cancer lesions

The above results suggest that tumor-associated nociceptors may initiate communications with incoming monocytes within the first few days of tumor development to shape an immunosuppressive milieu as early as day 7 when T-cells first begin to appear. Thus, we sought to determine at what time and for how long nociceptors are needed to promote tumor persistence. Since our *in vitro* studies had shown that M-MDSC differentiation does not depend on neuropeptides (**Figure S6E**), however, nociceptors needed to be alive (**Figure 6A**) and in direct contact with monocytes (**Figure 6C**) for M-MDSC formation to occur, we hypothesized that nociceptors must be electrically active to exert their effect on monocytes. To test this idea, we crossed NaV1.8*^Cre^* mice with *R26^LSL-Gi-DREADD-mCitrine^* animals^77^. In this strain, henceforth called NaV1.8*^Gi-DREADD^*, nociceptors expressed an inhibitory Designer Receptors Exclusively Activated by Designer Drugs (DREADD), a Gα_i/o_-linked GPCR that can be activated by an otherwise biologically inert ligand, C21 (compound 21), to induce hyperpolarization of neurons and inhibit their activity^78^. In dorsal root ganglia of NaV1.8*^Gi-DREADD^* mice, the mCitrine reporter, which indicates DREADD expression, was restricted to a subset of neurons and absent in Cre-negative littermates, confirming the specificity of Gi-DREADD expression (**Figure S7I**).

To ask whether the electrical activity of nociceptors is needed for BC formation, we subjected NaV1.8*^Gi-DREADD^* mice and littermate controls to daily C21 injections and instilled MB49 cells in their bladders. Strikingly, while all control mice developed tumors, the tumor incidence was significantly decreased in C21-treated NaV1.8*^Gi-DREADD^* mice (**Figure 7C**). Thus, sustained inhibition of electrical activity in nociceptors phenocopied the effect of nociceptor ablation, indicating that nociceptors are active participants in the process of bladder tumorigenesis. Finally, to clarify at which stage of tumor growth nociceptors are required, we divided tumor-implanted NaV1.8*^Gi-DREADD^*mice into two cohorts and treated them with daily injections of C21 either during the early (days 0-6) or late (days 7-13) stages of tumorigenesis. While all mice in the late treatment group developed tumors, mice that had received early C21 treatment exhibited a decreased tumor incidence and size, albeit not to the extent observed in nociceptor-ablated animals or in NaV1.8*^Gi-DREADD^* mice that received continuous C21 treatment (**Figure 7D**).

## Discussion

Here, we demonstrate that a three-way communication between tumor cells, nociceptors, and the immune system is critical in shaping the TME of urinary bladder carcinoma. Our results indicate that tumor cells invading the bladder wall recruit classical monocytes while simultaneously releasing neurotrophic factors that promote axon sprouting of local nociceptors. Electrical activity of nociceptors during the early stages of BC growth is pivotal to drive the differentiation of newly recruited monocytes into M-MDSCs to create an immunosuppressive microenvironment. At the same time, drainage of tumor antigens to local lymph nodes generates a population of tumor-specific CTLs. However, owing to their need to expand and differentiate from naive precursors and then traffic from lymph nodes to the bladder^79^, substantial numbers of CTLs appear in tumor-bearing bladders only ∼one week after tumor challenge, at which time M-MDSCs are already abundant and effectively prevent the newly arrived CTLs from eradicating most tumors (**Figure 8)**. Depletion of either monocytes or nociceptors or inhibition of nociceptor activity during this vulnerable period interrupted this process and rendered tumors highly susceptible to CTL-mediated rejection.

**Figure 8.**
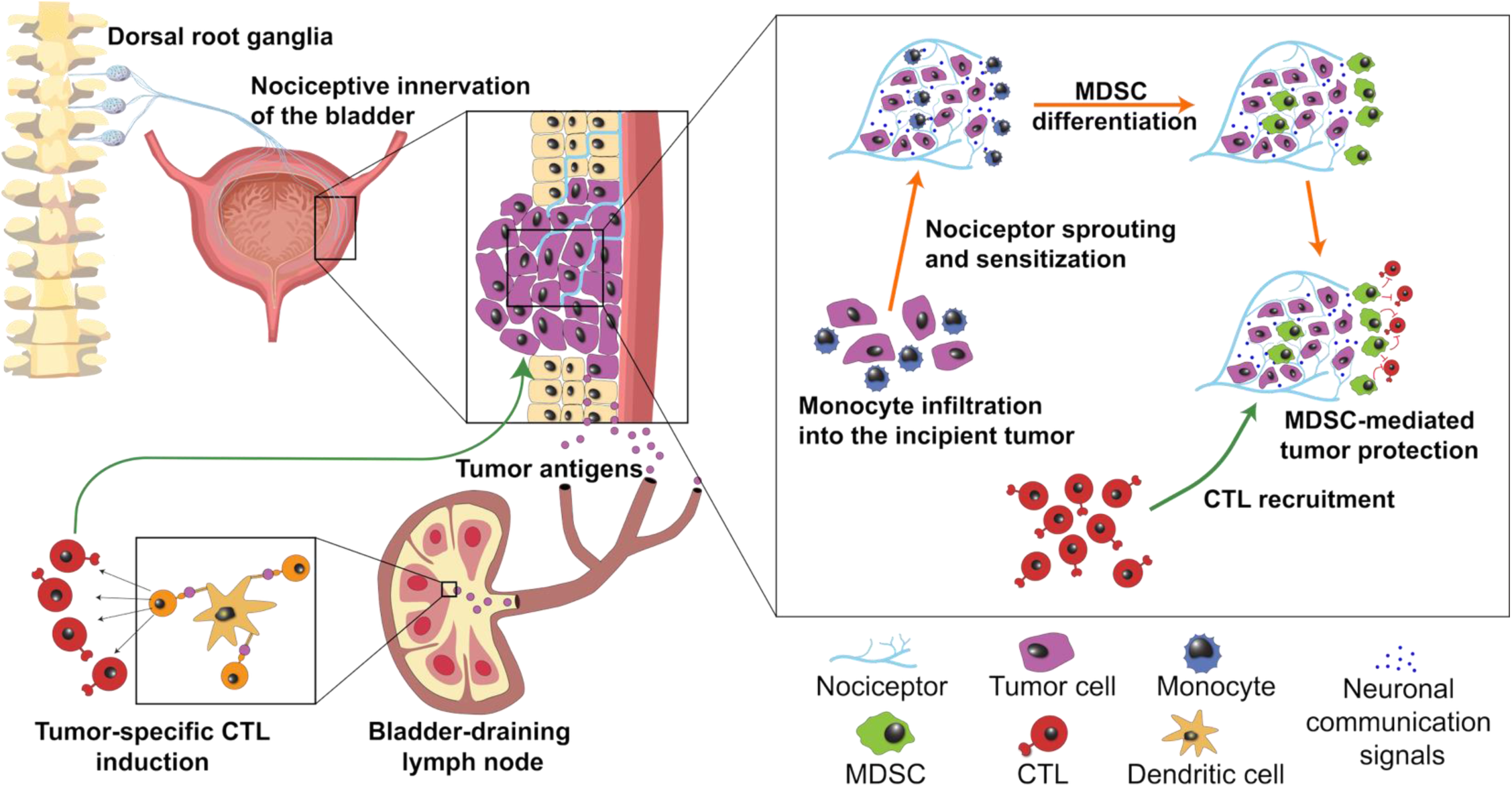
Schematic depiction of the three-way communication among bladder carcinoma, nociceptors, and monocytes.

### Nociceptors in bladder cancer

Our findings have potential clinical implications for BC therapy. Despite recent advances in the treatment of malignant diseases, BC remains a significant unmet clinical need. The standard of care for high-risk BC has not changed in 40 years and primarily consists of intravesicular instillation of Bacillus Calmette-Guerin (BCG)^80^. Although BCG therapy is initially highly efficacious, disease recurrence is common, with 1- and 5-year recurrence rates of 15–60% and 31–78%, respectively^81^. In light of this clinical experience, novel therapeutic approaches, such as modulators of the TME, are needed. In fact, several observations point to MDSCs as potential therapeutic targets^82^. For example, MDSC depletion has shown promise in mouse models of BC^83,84^, and the abundance of MDSCs within tumors is inversely correlated with the prognosis of BC patients^85^. Nonetheless, it has been unclear when and how the precursors of MDSCs are recruited to tumors, nor do we understand the mechanisms of intratumoral MDSC differentiation.

Our current results show that tumor-innervating nociceptors are highly effective at driving the rapid differentiation of inflammatory Ly6C^Hi^ monocytes into immunosuppressive cells with phenotypic and functional characteristics of M-MDSCs. It must be cautioned that the mere inhibition of nociceptors alone was not curative during later stages of the disease, likely owing to the already established presence of an immunosuppressive TME resulting in the appearance of ‘cold’ tumors where CTL are effectively excluded from accessing tumor cells. Nonetheless, this does not rule out the possibility that nociceptor targeting could provide a clinical benefit when combined with existing chemotherapy and/or immunotherapy regimens such as BCG or immune checkpoint blockade. Additionally, targeting nociceptors (or monocytes/MDSCs) might help tackle another long-standing unmet need in the treatment of BC: intraluminal seeding. Owing to their anatomic location, malignant cells that are discharged from the primary tumor into the bladder lumen can establish new lesions in distal, previously healthy parts of the organ^86^. As intraluminal tumor cell shedding can occur at any point during tumor growth, nociceptor inhibition could potentially curtail tumor spreading. In addition, a nociceptor targeted therapy might also be relevant during minimally invasive surgical treatment of BC. Transurethral resection of bladder tumors (TURBT), whereby abnormal tissue is excised from the bladder wall by cystoscopy, remains the standard of surgical care for BC. However, in the course of this procedure, resected tumor fragments often remain in the luminal cavity and surgeons rely on micturition for their expulsion. Re-implantation of resected primary tumor fragments can result in the formation of new lesions and disease recurrence^86^. Since the process through which MB49 cells are implanted in mice represents a model of intraluminal seeding^87^, it is tempting to speculate that pharmacological inhibition of bladder nociceptors during and after TURBT might improve outcomes by decreasing the rate of tumor recurrence.

### Immune cell subset- and tumor type-specific effects of nociceptors

Recent work by several groups has identified critical tumor-promoting roles for nociceptors in other malignancies^19–24^. In most settings, the nociceptor-derived mechanistic drivers of tumor progression were neuropeptides, particularly CGRP ^19–21,23^ and substance P ^24^, which contributed to T cell exhaustion or exerted direct growth-promoting effects on cancer cells. By contrast, in the present study, nociceptors exerted a protective effect on tumors by a fundamentally distinct mechanism that did not rely on neuropeptides. Non-neuropeptide-mediated modalities of communication between nociceptors and immune cells have been reported before^9,88,89^. In the orthotopic BC model used here, the pro-tumorigenic nociceptor effect resulted from a profound modulation of the TME and required that nociceptors were electrically active to promote MDSC differentiation of inflammatory monocytes.

Remarkably, although the presence of nociceptors in cocultures also impacted the transcriptional profiles of the patrolling monocyte subset as well as that of neutrophils, these myeloid cells did not acquire a full-fledged MDSC phenotype in cocultures. Similarly, we have shown recently that nociceptors use at least three distinct communication modalities, including electrical activity, to regulate the function of DCs^9^, but the net impact of nociceptors on DCs is pro-inflammatory, rather than immunosuppressive^1,2^. Thus, while nociceptors can apparently modulate most, if not all, myeloid leukocyte subsets^49^, their effect on classical monocytes described here appears to be unique.

### Tumor location-specific effects of nociceptors

Beyond their immune cell subset and tumor type specific impact, the role of nociceptors in tumor development also appears to vary with anatomic context. In the MB49 BC model used here, nociceptors were critical for orthotopic tumor formation in the bladder, but not in the subcutis. Although SQ implanted MB49 tumors are often considered a surrogate model of BC^29,90^, our findings indicate that the developmental trajectories of the TME within ectopic vs. orthotopic MB49 lesions differ substantially. Specifically, unlike in the bladder, MB49 tumors in the subcutis exhibited an early abundance of cells with the phenotypic markers of neutrophils (Ly6C^Lo^ Ly6G^+^ CD11b^+^), and did not appear to receive nociceptive innervation. While the reasons for these differences remain to be further explored, we note that in the orthotopic setting, only a handful of tumor cells invade the urothelial barrier to form one or more subepithelial infiltrating lesions that are densely innervated throughout the tumor’s development. By contrast, the experimental protocol for the SQ model requires the injection of a large number of suspended tumor cells that are deposited under the skin and then grow concentrically as a compact nodular tumor embedded in a dense capsule-like layer of connective tissue. Histologically, nociceptors were abundant in the immediate vicinity of SQ MB49 tumors but rarely seen to penetrate the capsule. It is possible that this rather artificial approach to induce SQ tumors by local injection of a tumor cell suspension, whereby tumors displace rather than infiltrate the surrounding host tissue, may be less conducive to sensory innervation. However, in contrast to SQ MB49 tumors, a recent study has demonstrated that SQ B16 melanoma, which was induced using a similar cell injection protocol, promoted nociceptive innervation^19^. It is thus possible that the lack of innervation observed in SQ MB49 tumors may not be inherent to the subcutis *per se*, but may rather result from differences in the way urothelial tumor cells vs. melanoma cells interact with the cutaneous environment.

An additional potentially relevant difference between orthotopic and SQ MB49 tumors was the delayed kinetics of PMN-MDSC recruitment in the former. The exact role of this myeloid population within the TME remains to be further defined, however, it is conceivable that their early appearance in SQ tumors is sufficient to counteract anti-tumor CTL responses, so in this setting any nociceptor-dependent M-MDSC differentiation may be redundant.

These anatomic distinctions raise questions about the contribution of nociceptors to metastatic disease, the main driver of cancer mortality^91^. About half of all patients with muscle-invasive bladder cancer develop metastases within 2 years, and metastatic bladder cancer has a 5-year survival rate of 5% ^92^. The most commonly affected organs in metastatic BC are lymph nodes (70%), bone (47%), lung (37%), liver (26%), and peritoneum (16%), followed by pleura (11%), adrenal glands (7%), and brain (5%)^93^. Whether, when, and to what extent nociceptors and nociceptor-induced M-MDSCs contribute to lesion formation in these secondary target tissues will require further exploration.

### Nociceptive activity in the absence of sensory perception of pain

The precise cellular and molecular mechanisms by which electrical activity in nociceptors compels classical monocytes to proliferate and assume an M-MDSC phenotype will also require further elucidation. Notwithstanding, our chemogenetic experiments imply that nociceptive electrical activity during the early stages of tumorigenesis is essential for the establishment of bladder tumors. Firing nociceptors are traditionally thought to send afferent signals to the CNS that are perceived as pain or itch. Yet, most patients with BC initially present with painless hematuria^94^. Accordingly, mice with MB49 orthotopic tumors do not display morphine-seeking behavior, indicating an analogous lack of sensory experience^95^. Similarly, although nociceptors are involved in the development of melanoma, pain or itch in the primary tumor are uncommon symptoms^96^. Moreover, recent work has shown that spleen-innervating nociceptors promote antibody responses during flu infection and after vaccination^12^ even though neither condition is generally associated with splenic discomfort or pain. Thus, our observations in BC add to a growing list of seemingly indolent conditions in which nociceptor activity, nonetheless, exerts control over the immune system. At least in some settings, the immunoregulatory and sensory functions of nociceptors in the periphery can apparently be uncoupled from their afferent functions in the CNS, raising the possibility that nociceptors may be involved in other pathologies that are not associated with overt pain.

### Limitations of the study

In all our experiments, we ablated or inhibited nociceptors systemically. Consequently, we cannot exclude the possibility that nociceptors outside the bladder might also play relevant roles. Indeed, nociceptors can affect antigen trafficking to lymph nodes^97^ as well as the ability of dendritic cells to elicit T-cell responses^9^. The activation of tumor antigen-specific T-cells we observed in RTX-treated animals demonstrates that nociceptors are not essential to this process; however, we cannot rule out that they may play more subtle roles. Additionally, our co-culture experiments revealed that nociceptors, in a neuropeptide-independent manner, induce the acquisition of an immunosuppressive M-MDSC-like phenotype in primary monocytes by activating, at least in part, the Sp1 and Hif1α signaling pathways. Both Sp1 and Hif1α pathways regulate numerous cellular processes and are themselves regulated by many signaling molecules, suggesting that a broader approach, which is beyond the scope of the current study, will be necessary to establish if both pathways are activated by a single communication event or if multiple mechanisms are involved, and to fully unravel how the MDSC transition is mediated at a molecular level.

## Materials and Methods

### Mice

C57BL/6J (JAX stock no. 000664), *Calca^Cre-GFP^* (JAX stock no. 033168)^69^, and *R26^LSL-Gi-DREADD^*(JAX stock no. 026219)^77^ mice were purchased from the Jackson Laboratory. *Scn10a^Cre^* ^98^*, Scna10a^Cre^ R26^Ai^*^14^ ^38^*, and Scn10a^Cre^Ccl2-mCherry^fl/fl^* ^9^ were all described previously and were bred at the Harvard Medical School animal facility. All animal experiments were performed in accordance with national and institutional guidelines and were approved by the institutional animal care and use committee and the Committee on Microbiological Safety (COMS) of Harvard Medical School.

### Cells

MB49 cells were purchased from EMD Millipore and cultured in DMEM medium (Corning) supplemented with 10% FCS (Gemini) and penicillin-streptomycin. For *in vivo* experiments, the cells were washed twice in PBS and kept on ice until implanted.

MB49-eGFP cells were generated by lentiviral transduction of the parental MB49 line as described previously.^99^ Briefly, 90% confluent Lenti-X 293T cells (Takara Bio) grown in a 10 cm dish were transfected with 15 μg of pLenti PGK GFP Puro (Addgene plasmid #19070), 2.4 μg of pRSV-rev (Addgene plasmid #12253), 4 μg of pMDLg/pRRE (Addgene plasmid #12251), and 1.8 μg of pMD2.G (Addgene plasmid #12259) using Lipofectamine 3000 (Thermo Fisher). After 24 hours, the medium was replaced with 5 ml of complete DMEM. Virus-containing medium was harvested 24 hours later, clarified by centrifugation and filtration through a 0.45 μm filter, and applied onto 70% confluent MB49 cells. After 24 hours, the medium was replaced with fresh complete DMEM, and, after a further 24 hours, 3μg/ml puromycin was used to select the transduced MB49 cells.

SIINFEKL-MB49 cells were generated by cloning the mScarlet-SIINFEKL expression cassette from medSIIN (Addgene plasmid #185664) into pLenti-PGK GFP Puro (w509-5) (Addgene plasmid #19070), replacing the GFP sequence to generate the new pLenti-PGK-mScarlet-SIIN vector. The construct does not include a Kozak sequence upstream of the expression cassette, resulting in only moderate SIINFEKL expression. MB49 cells were stably transduced with lentivirus packaged from the pLenti-PGK-mScarlet-SIIN vector, and transduced cells were subsequently sorted based on mScarlet expression.

*Ngf-*KO MB49 cells were generated by CRISPR-Cas9-mediated deletion using a gRNA (TGGGTGCTGAACAGCACACG) targeting exon 4 as described previously ^100^. Briefly, sgRNA (Synthego) was precomplexed with Cas9 protein (IDT) at room temperature for 10 minutes and nucleofected into parental MB49 cells with SE-DS150 buffer-pulse combination using the 4D Nucleofector (Lonza). Limiting dilution was performed to obtain monoclonal cell populations, and expression of NGF was assessed by Western blot.

*Ngf* reconstitution of *Ngf*-KO MB49 cells was performed by replacing the GFP cassette in the pLenti PGK GFP Puro plasmid (Addgene plasmid #19070) with *Ngf* cDNA amplified from parental MB49 cells. *Ngf*-KO MB49 cells were stably transduced with lentivirus packaged from the pLenti PGK Ngf Puro plasmid and selected in puromycin as described above for MB49-eGFP cells.

### SDS-PAGE and Immunoblot analysis

2x10^6^ MB49 cells were harvested, washed once in PBS, and lysed on ice for 30 minutes in 1% NP40 in PBS containing protease inhibitor cocktail (Thermo Scientific). Lysates were clarified by centrifugation and prepared for SDS-polyacrylamide gel electrophoresis (SDS-PAGE) in 6× Laemmli buffer containing 60% v/v glycerol, 150 mg/ml SDS, and 0.75 mg/ml bromophenol blue in 75 mM Tris–HCl, pH 6.8, in the presence of 100 mM dithiothreitol. Separation was performed using 4 to 20% Mini-PROTEAN TGX precast gels (Bio-Rad). Proteins were transferred onto an Immobilon P membrane (Merck Millipore), and the membrane was blocked in 5% milk in PBS + 0.05% Tween-20. E7O4E rabbit anti-NGF (Cell Signaling) and 13E5 rabbit anti-β-actin (Abcam) antibodies were used to detect the respective proteins. Signal was revealed using Luminata Forte HRP substrate (Merck Millipore) on the ChemiDoc MP imager (Bio-Rad).

### Staining reagents

Arg-1, B220, β3-tubulin, CCR2, CD3ε, CD4, CD8α, CD8β, CD11b, CD11c, CD19, CD29, CD34, CD39, CD44, CD45, CD64, CD84, CX3CR1, E-Cadherin, Galectin 3, Gr-1, Hif1α, ICOS, Ki67, Laminin, Ly6C, Ly6G, mCherry, MHC-II, NK1.1, PD-L1, Siglec F, ST2, TCRβ, TCRψο, Thy1, SIINFEKL-tetramers

### Tumor experiments

Orthotopic MB49 tumors were implanted as described previously.^31^ Briefly, 6-12-week-old female mice were anesthetized by the combination of ketamine and xylazine, immobilized in the supine position, and had their bladders voided by applying gentle pressure on the abdomen. Lubricated PE10 polyethylene tubing (BD Intramedic) connected to a syringe with a 26-gauge needle was transurethrally inserted into the bladder, fixed in place, and 100μl of 100μg/ml poly-D-Lysine (Millipore Sigma) was instilled into the bladder. After 1 hour, poly-D-Lysine was evacuated by gentle pressure on the abdomen, and 1x10^5^ MB49 cells in 50 μl of PBS were instilled. After 1 hour, the catheters were removed, the bladders were voided by gentle pressure on the abdomen, and the mice were allowed to emerge from anesthesia. Mice were euthanized and their bladders analyzed on day 14 after tumor implantation unless indicated otherwise.

For DREADD experiments, tumor-implanted mice were injected intraperitoneally with 3mg/kg of C21 (Tocris) in 150 μl of PBS according to the regimens outlined in Fig. 4C and D.

For monocyte depletion experiments, the MC21 anti-CCR2 antibody was used as described previously^35,36^. Briefly, 20 μg of MC21 or isotype control (IgG2b,κ) antibody was injected daily (i.p.), starting from day 4 post-tumor implantation. Monocyte presence in the blood was assessed on days 7 and 9, and mice with incomplete monocyte depletion were excluded from the experiment.

Ectopic Sub-Q MB49 tumors were implanted as described previously.^29^ Briefly, 6-12-week-old female mice were anesthetized by isoflurane, and 1x10^4^ or 1x10^5^ MB49 cells in PBS were injected into the scruff. Mice were euthanized on day 14 after tumor implantation unless indicated otherwise.

### In vivo nociceptor ablation

RTX-mediated nociceptor ablation was performed as previously described.^11^ Briefly, four-week-old mice were injected sub-Q in the scruff on three consecutive days with 30, 70, and 100 mg/kg of RTX (AdipoGen Life Sciences) or vehicle control (DMSO) and allowed to age at least 6 weeks before being used in experiments. Functional denervation was assessed by the “tail flick” assay.

### Flow cytometry

Mouse urinary bladders and lymph nodes were harvested, placed on ice, cut with scissors into small pieces, and digested for 1 hour at 37°C with constant shaking in RPMI1640 medium containing 100μg/ml DNAse (Sigma Aldrich) and 62.5μg/ml TM Liberase (Roche). Single-cell suspensions were generated in the GentleMACS M tubes on a GentleMACS tissue dissociator (Miltenyi Biotec) and further homogenized by mechanical disruption on 50 μm cell strainers. Single-cell suspensions were kept on ice, dead cells were labeled using the LIVE/DEAD Fixable Near-IR Dead Cell Stain Kit (Thermo Fisher), Fc receptors were blocked with the anti-CD16/32 antibody, and surface markers were stained with the appropriate fluorescent antibodies in FACS buffer (1% FBS, 5 mM EDTA, and 0.1% NaN_3_). Where necessary, intracellular staining was performed using the BD Cytofix/Cytoperm Fixation/Permeabilization Kit (BD Biosciences) as per the manufacturer’s instructions. Data acquisition was performed on a Cytoflex S (Beckman) flow cytometer, and the data were analyzed in FlowJo v10.9.0 software. Individual immune cell subsets were identified as shown in Fig. S1E.

*In vitro* cultures were harvested in 5 mM EDTA in PBS. Dead cells were labeled using the LIVE/DEAD Fixable Near-IR Dead Cell Stain Kit (Thermo Fisher), Fc receptors were blocked with the anti-CD16/32 antibody, and surface markers were stained with the appropriate fluorescent antibodies in FACS buffer (1% FBS, 5 mM EDTA, and 0.1% NaN_3_ in PBS). Where necessary, intracellular staining was performed using the BD Cytofix/Cytoperm Fixation/Permeabilization Kit (BD Biosciences) as per the manufacturer’s instructions. Data acquisition was performed on a Cytoflex S (Beckman) flow cytometer, and the data were analyzed in FlowJo v10.9.0 software.

### Bladder imaging

For confocal imaging of bladder preparations, the mice were euthanized, and their bladders were evacuated by applying pressure to the abdomen. An incision was made in the abdominal wall, and the bladder was exposed. Lubricated polyethylene tubing (BD Intramedic) connected to a syringe with a 26-gauge needle was transurethrally inserted into the bladder, and 4% paraformaldehyde (PFA) was instilled. The bladder was maintained in an inflated state and irrigated with PFA from the outside for 20 minutes. For whole-mount imaging, the fixed bladder was dissected, cut in half, permeabilized in 0.3% Tween in PBS, blocked in 3% BSA, and stained with appropriate antibodies. For cryo-sectioning, PFA was removed from the bladder and replaced with 50% OCT (Sakura) in 30% sucrose. Tubing was rapidly removed, and the inflated bladder was submerged in OCT and flash-frozen on dry ice. Cryo-sectioning was performed on a Cryostar NX70 cryostat (Thermo Scientific), and 50-100 μm thick sections were generated, permeabilized in 0.3% Tween in PBS, blocked in 3% BSA, and stained with appropriate antibodies. For optical clarification, the iDISCO protocol^39^ was used with minor modifications. Briefly, whole bladders were fixed overnight in 4% PFA, washed for 1 hour in 50% MetOH, 1 hour in 100% MetOH, 1 hour in 50% MetOH, 45 minutes in PBS, 90 minutes in 0.2% Tween in PBS, and incubated overnight in PBS containing 0.2% Tween, 20% DMSO, and 0.3 M Glycine. Samples were blocked for 24 hours in PBS containing 0.2% Tween, 10% DMSO, and anti-CD16/32 antibodies (Fc-block), washed twice for 1 hour in PBS containing 10 µg/ml heparin, and stained for 5 days in PBS containing 5% DMSO, 10 µg/ml heparin, Fc-block, and appropriate antibodies. Following staining, the samples were washed 6 times for 1 hour and incubated overnight in PBS with 10 µg/ml heparin. Finally, the samples were dehydrated by a 1-hour treatment with 50% MetOH and 3 subsequent treatments with 100% MetOH, and either stored in 100% MetOH at -20°C or cleared overnight in a 1:1 mixture of benzyl alcohol and benzyl benzoate.

All imaging was performed on the Olympus IX83 inverted single-point laser scanning confocal microscope with Uplan S Apo 4×/0.16 air, UPlan S Apo 10×/0.40 air, UPlan X Apo 20×/0.8 air, or UPlan X Apo 60×/1.42 oil objectives.

For H&E analysis, uninflated mouse bladders were dissected and fixed in formalin, embedded in paraffin, sectioned, and stained with hematoxylin and eosin. Images were acquired on the Olympus VS200 Slide Scanner with UPlan X Apo 10×/0.40 air objective.

### Monocyte isolation

Spleens of mice were harvested, cut with scissors into small pieces, and digested for 1 hour at 37°C with constant shaking in RPMI1640 medium containing 100μg/ml DNAse (Sigma Aldrich) and 62.5 μg/ml TM Liberase (Roche). Single-cell suspensions were generated by mechanical disruption on 50 μm cell strainers in MACS buffer (1% FBS, 2 mM EDTA in PBS). Red blood cells were lysed in ACK buffer, dead cells were labeled using the LIVE/DEAD Fixable Near-IR Dead Cell Stain Kit (Thermo Fisher), Fc receptors were blocked with the anti-CD16/32 antibody, and cells were stained with fluorescent antibodies and FACS-sorted on a MoFlo Astrios EQ FACS-sorter (Beckman). Classical monocytes were identified as CD3ε^-^ TCRβ^-^ CD19^-^ B220^-^Ly6G^-^ CD11b^+^ Ly6C^Hi^, non-classical as CD3ε^-^ TCRβ^-^ CD19^-^ B220^-^ Ly6G^-^ CD11b^+^ Ly6C^Lo^ CX3CR1^+^.

### In vitro co-culture experiments

Nociceptor cultures were prepared and maintained as described previously.^9,101^ Immediately before the monocyte co-culture, the nociceptor medium was replaced with 100 μl of fresh B27-supplemented Neurobasal medium (Gibco), and FACS-purified monocytes were added in 100 μl of RPMI1640 medium with 10% FCS. Cells were co-cultured overnight unless otherwise indicated, and analyzed the next morning. For culture treatments, LPS (Thermo) was used at 100ng/ml, 8-Br-cAMP (Sigma Aldrich) at 1mM, forskolin (Selleckchem) at 50μM, mithramycin A (Cayman Chemical) at 50 and 500nM, echinomycin (Abcam) at 5nM, DMOG (Cayman Chemical) at 100μM, and all neuropeptides (Phoenix Pharmaceuticals) at 1μM.

For axon outgrowth assays, nociceptors were purified as described previously and cultured overnight in B27-supplemented Neurobasal medium with penicillin-streptomycin but without NGF and cytosine β-D-arabinofuranoside in the presence or absence of 10^4^ MB49 cells.

For CGRP release assays, nociceptor cultures were prepared and maintained as described previously and cultured for 7 days in B27-supplemented Neurobasal medium with penicillin-streptomycin, 5 mM cytosine β-D-arabinofuranoside (Sigma-Aldrich), and 25 ng/ml mouse recombinant NGF (R&D Systems). The culture medium was refreshed on day 4. On day 7, the medium was replaced with a 1:1 mixture of DMEM medium (Corning) supplemented with 10% FCS (Gemini), and B27-supplemented Neurobasal medium with penicillin-streptomycin but without NGF and cytosine β-D-arabinofuranoside and 10^4^ MB49 cells were added. After 24 hours of co-culture, 1μM capsaicin was added for 2 hours.

### In vitro T-cell proliferation experiments

Flt3L-induced bone marrow dendritic cells (BM-DCs) were pulsed with 100μg/ml ovalbumin for 2 hours at 37°C, washed twice, and cocultured with MACS-purified, cell trace violet-loaded, ovalbumin-specific CD8 T-cells (OT-I), and FACS-purified tumor-derived M-MDSCs, PMN-MDSCs, or splenic monocytes or neutrophils at a 1:1:1 ratio. T-cell proliferation was assessed by flow cytometry on day 3.

### Cytospins

Tumor-derived M-MDSCs and PMN-MDSCs, splenic and healthy bladder monocytes, and bone marrow and splenic neutrophils were FACS-purified and deposited onto a slide by 2-minute centrifugation at 300 RPM in a Cytopro cytocentrifuge (ELITech Biomedical Systems). Cells were fixed, stained with hematoxylin and eosin or toluidine blue, and imaged on the Olympus VS200 Slide Scanner with UPlan X Apo 60×/1.42 oil objective.

### RNA sequencing

Monocytes or neutrophils cultured in the presence or absence of nociceptors overnight, or primary splenic monocytes and tumor-derived M-MDSCs were FACS-purified on a MoFlo Astrios EQ FACS-sorter (Beckman). One thousand cells of each population were sorted directly into 5 μl of lysis buffer (TCL Buffer from Qiagen with 1% β-mercaptoethanol), and Smart-seq2 libraries were prepared as previously described.^102,103^ Briefly, total RNA was captured and purified on RNAClean XP beads (Beckman Coulter). Polyadenylated mRNA was then selected using an anchored oligo(dT) primer (5′ AAGCAGTGGTATCAACGCAGAGTACT30VN 3′) and converted to cDNA via reverse transcription. First strand cDNA was subjected to limited PCR amplification followed by transposon-based fragmentation using the Nextera XT DNA Library Preparation Kit (Illumina). Samples were PCR amplified for 18 cycles using barcoded primers such that each sample carried a specific combination of eight-base Illumina P5 and P7 barcodes and pooled together prior to sequencing. Paired-end sequencing was performed on an Illumina NextSeq500 using 2 x 25bp reads.

### RNAseq data analysis

Reads were aligned to the mouse genome (GENCODE GRCm38/mm10 primary assembly and gene annotations vM16; https://www.gencodegenes.org/mouse_releases/16.html) with STAR 2.5.4a (https://github.com/alexdobin/STAR/releases). The ribosomal RNA gene annotations were removed from GTF (General Transfer Format) file. The gene-level quantification was calculated by featureCounts (http://subread.sourceforge.net/). Raw reads count tables were normalized in iDEP2.0 (EdgeR (Log2(CPM+1))).^104^ Only genes with a minimal 50 counts per million (CPM) expression in at least two libraries were selected for further analysis to exclude genes with little to no expression across all conditions. iDEP2.0^104^ was used to identify differentially expressed genes using DESeq2 with FC ≥ 2 and FDR 0.1 cutoffs, and to perform GO term enrichment analysis.^50,51^ Metascape^105^ was used to perform TRRUST analysis.^70^ GSEA v4.0.1 software was used to perform GSEA^63,64^ using manually generated lists of differentially regulated genes between M-MDSCs and monocytes and between monocytes cultured in the presence and absence of nociceptors. Heat maps were generated using Morpheus (https://software.broadinstitute.org/morpheus).

### Image analysis

Image analyses and visualizations were performed in Fiji 2.14.0^106^, QuPath 0.5.1^107^, and Imaris 9.5.0. Quantification of sprouting nociceptor axons was performed in Fiji, using the WEKA Image Segmentation script version 3.3.4^108^. A classifier was trained to segment axons, neuronal cell bodies, and background by manually annotating multiple fields of view containing diverse morphological features to ensure robust generalization across samples. Accuracy was confirmed manually on a small sample. The classifier was used in a Lusca^109^ script to automate the analysis on a larger scale.

### Survival analysis

Survival analysis was performed using KMplotter (https://kmplot.com/analysis/) as described previously^110^. Briefly, all possible cutoff values between the upper and lower quartiles for each gene were considered, and the cutoff value with the lowest false discovery rate (FDR) was selected. Patients were stratified according to the selected cutoff into the high (upper quartile) and low (lower quartile) expressors, and Cox proportional hazards regression analysis was performed to calculate differential survival rates. Kaplan-Meier plots were generated based on the survival data. Median survival was calculated for survival curves that drop to or below the 50% mark.

## Acknowledgments

We thank members of the von Andrian lab for helpful advice and discussions. We thank the HMS Flow Cytometry and Microscopy Resources on the Northern Quad (MicRoN) cores, and the DF/HCC Rodent Histopathology core for technical assistance. This work was supported by the HMS Center for Immune Imaging, grants from the National Institutes of Health (R01AI155865, R01AI175379) to U.H.v.A, a CRI Irvington postdoctoral fellowship (CRI2453) to P.H., a Gene Lay postdoctoral fellowship to C.P., and an Erasmus+ Worldwide Mobility Grant (#26595) of the European Union to A.B.

## SUPPLEMENTARY FIGURE LEGENDS

**Supplementary Figure 1.**
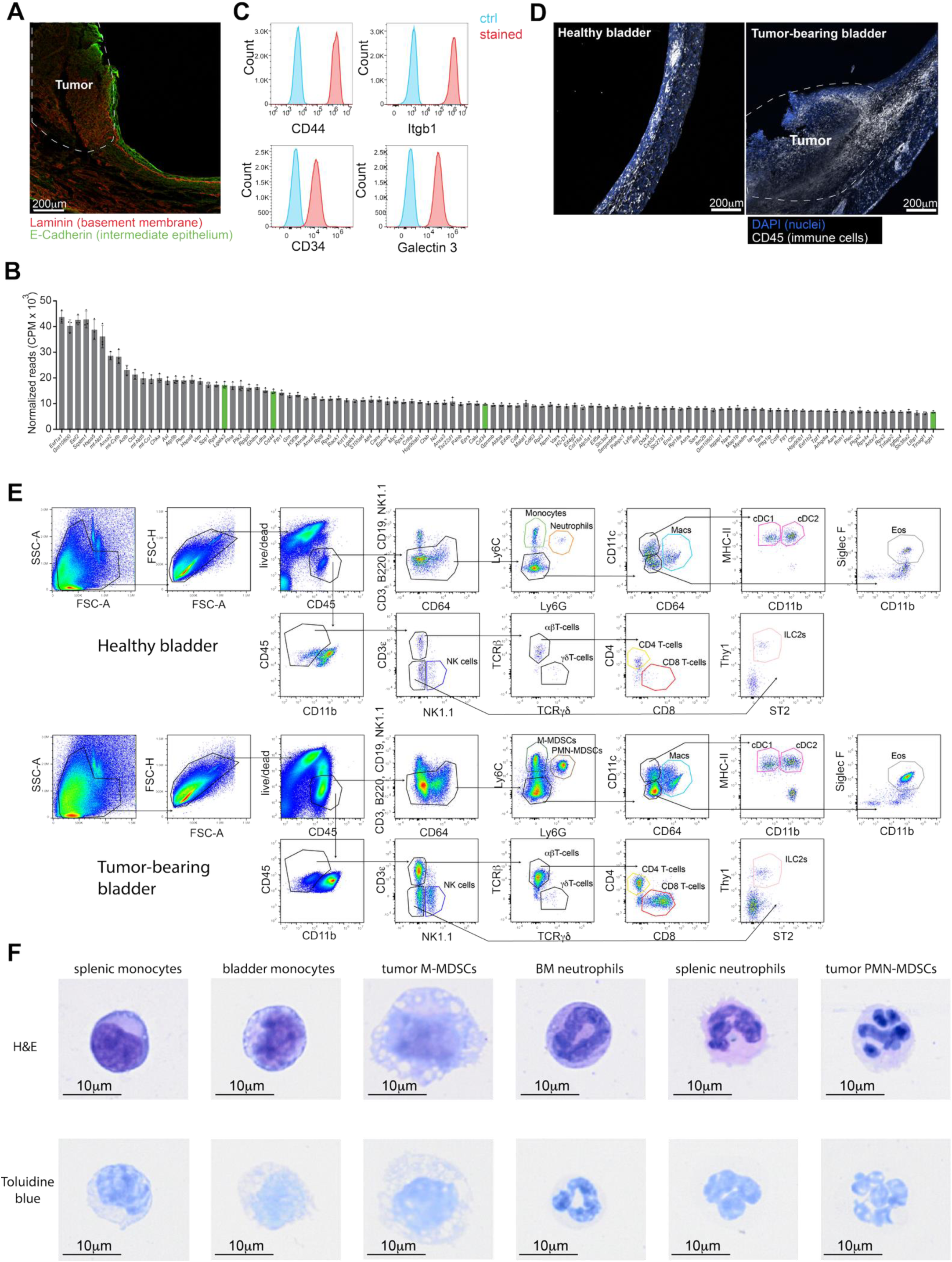
Phenotypic characterization of established MB49 bladder tumors and associated immune cell infiltrate. **(A)** The bladder of an MB49 tumor-bearing mouse was harvested on day 14 after tumor cell implantation, fixed, cryosectioned, stained with the indicated antibodies, and visualized by confocal microscopy. (**B**) MB49 cells were analyzed by RNAseq, and the most highly expressed genes are shown. Genes encoding surface molecules selected as candidate MB49 markers are highlighted in green. (**C**) MB49 cells were stained with the indicated antibodies and analyzed by flow cytometry. (**D)** Bladders of an MB49 tumor-bearing mouse and a healthy control were harvested on day 14 after tumor cell implantation, fixed, cryo-sectioned, stained with DAPI and anti-CD45 MAb, and visualized by confocal microscopy. (**E**) Gating strategy for the identification of immune cell types in healthy and MB49 tumor-bearing bladders. (**F**) FACS-purified MB49 tumor-derived M- and PMN-MDSCs, healthy splenic and bladder monocytes, and healthy bone marrow and splenic neutrophils were stained with H&E or toluidine blue and imaged using a slide-scanner microscope.

**Supplementary Figure 2.**
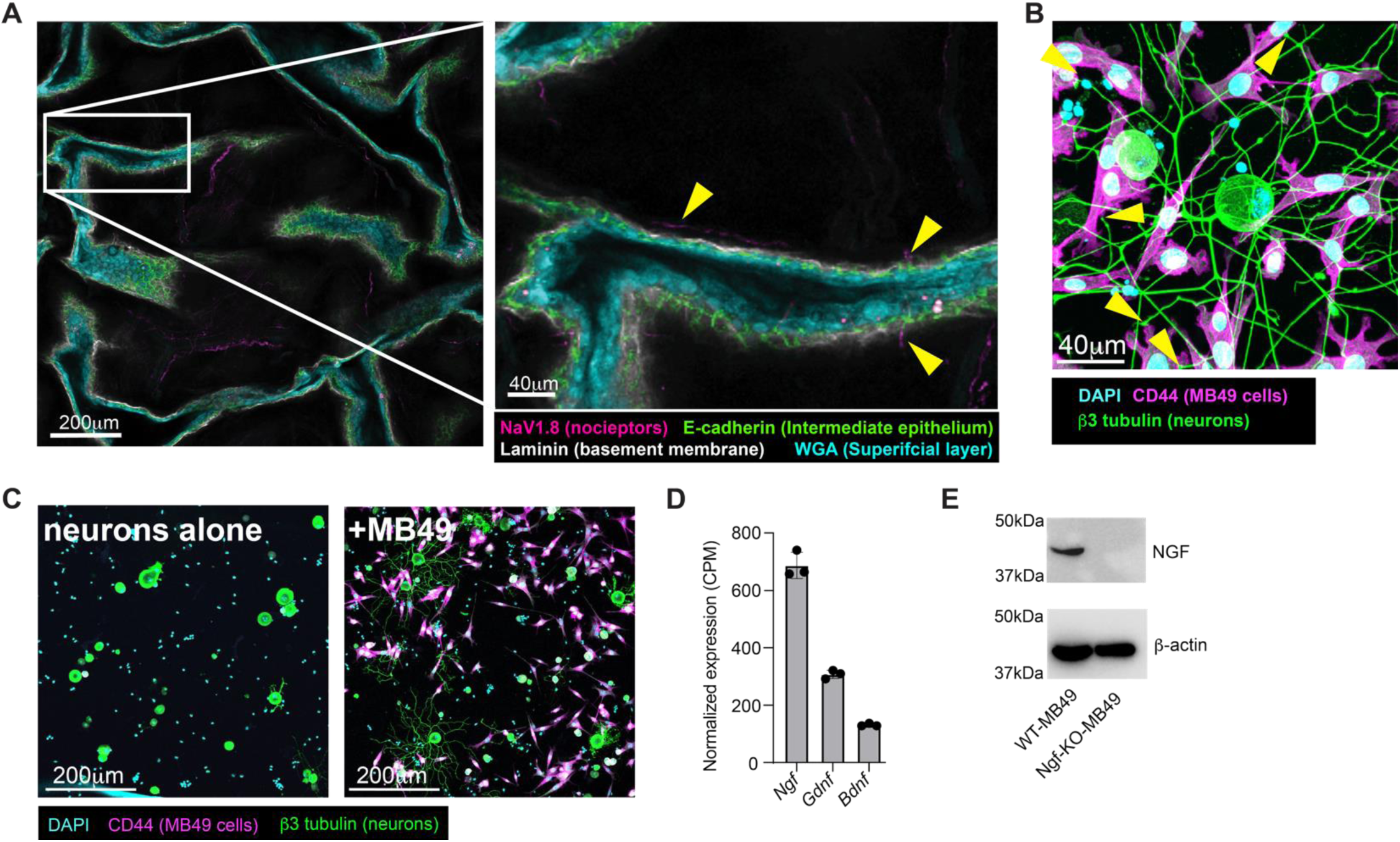
Interaction between nociceptors and MB49 cells. (**A**) A healthy mouse bladder was harvested, fixed, stained with the indicated antibodies, and imaged as a whole mount by confocal microscopy (see also **Suppl. Video 1**). A single optical section is shown, depicting the folds of deflated bladder tissue. Nociceptor fibers are highlighted by arrow heads. (**B**) MB49 cells were co-cultured with nociceptors for 3 days, fixed, stained with the indicated antibodies, and analyzed by confocal microscopy. (**C**) Freshly harvested, de-axonized nociceptors were cultured for 16 hours alone or in the presence of MB49 cells, fixed, and imaged by confocal microscopy. (**D**) MB49 cells were analyzed by RNAseq to assess gene expression of the indicated neurotropic factors. (**E**) Lysates from WT or *Ngf*-KO MB49 cells were analyzed by Western blot, probing for NGF or β-actin as a loading control.

**Supplementary Figure 3.**
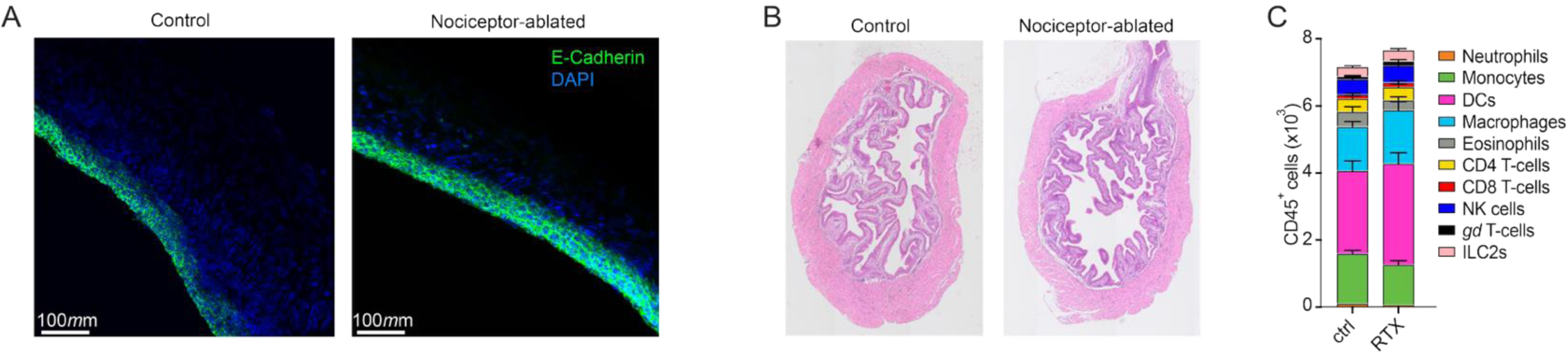
Healthy bladders remain grossly normal after nociceptor ablation. Healthy bladders of nociceptor-ablated and -competent mice were harvested and (**A**) fixed, cryo-sectioned, stained with the indicated antibodies, and visualized by confocal microscopy, (**B**) fixed, paraffin-sectioned, stained with H&E and visualized by brightfield microscopy, or (**C**), dissociated into single cell suspensions, and analyzed by flow cytometry (n=13-25, error bars indicate SEM).

**Supplementary Figure 4.**
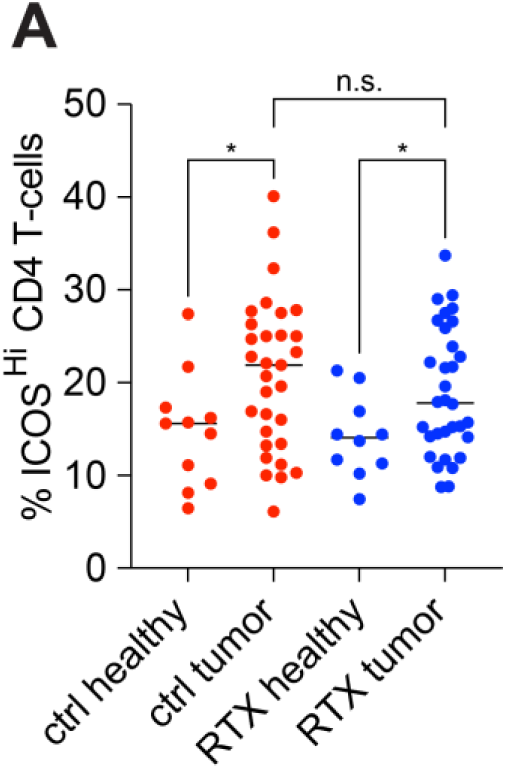
Activated CD4 T-cell in lymph nodes draining normal and tumor bearing bladders. Bladder draining lymph nodes of healthy or tumor-implanted nociceptor-ablated (RTX) and -competent (ctrl) mice were harvested on day 14 and analyzed by flow cytometry. Activated CD4^+^ T cell numbers were assessed by staining for ICOS and CD4. The data represent 10-12 healthy and 30-37 tumor-implanted animals per group. Two-way ANOVA with Tukey’s multiple comparisons test was used for statistical analysis. *p<0.05.

**Supplementary Figure 5.**
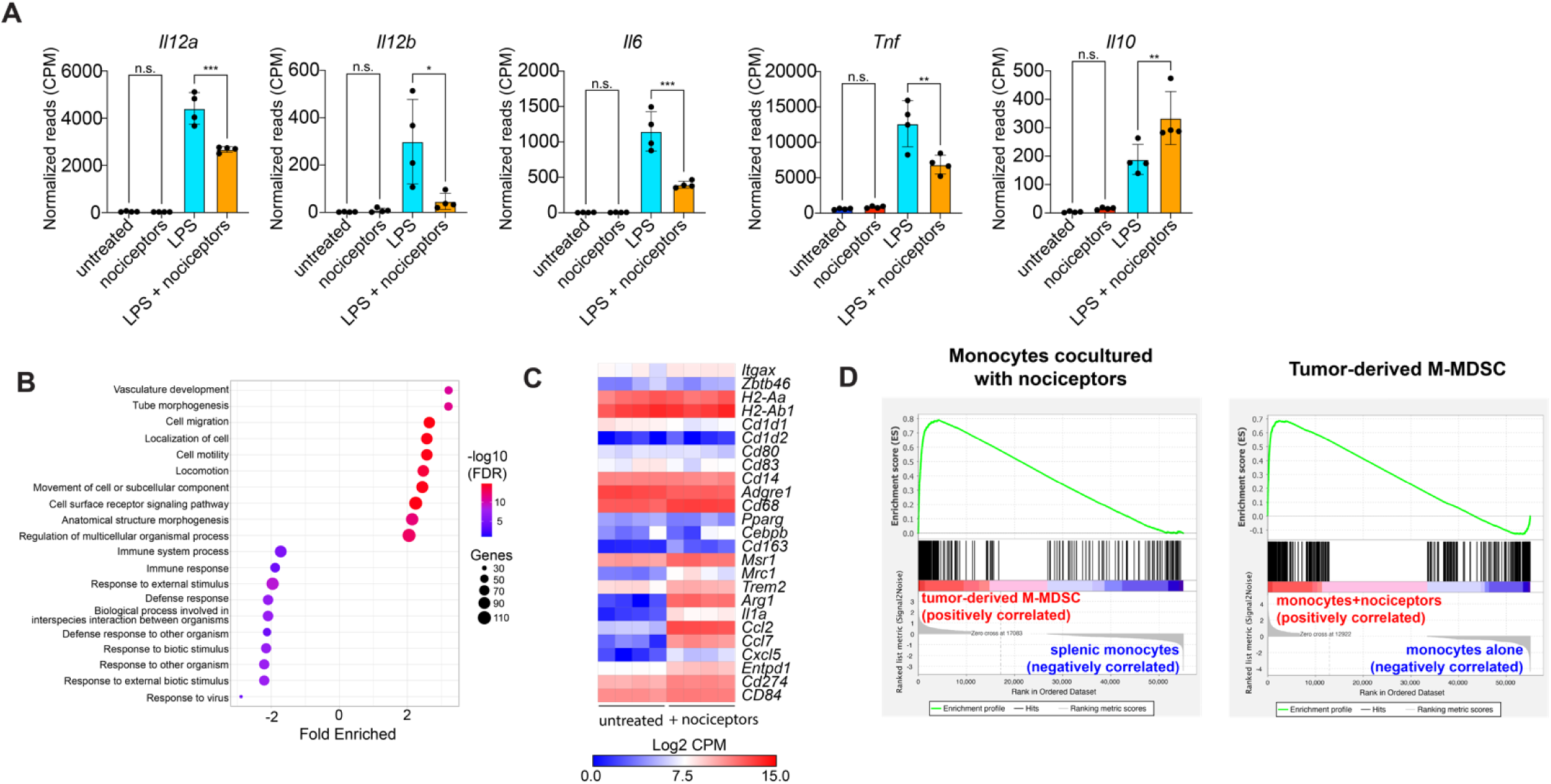
Nociceptor-induced changes in the transcriptome of monocytes. (**A-D**) FACS-purified Ly6C^Hi^ splenic monocytes were cultured alone or co-cultured with nociceptors in the presence or absence of LPS overnight, and analyzed by RNAseq. The expression of indicated cytokines (**A**), enrichment of GO terms comparing monocytes cultured alone and in the presence of nociceptors (**B**), and a heatmap illustrating the expression of indicated genes (**C**) are shown. (**D**) Gene-set enrichment of monocytes co-cultured with nociceptors, comparing them to primary tumor-derived M-MDSCs and primary splenic monocytes (left), and tumor-derived M-MDSCs, comparing them to monocytes co-cultured with nociceptors and cultured in isolation (right).

**Supplementary Figure 6.**
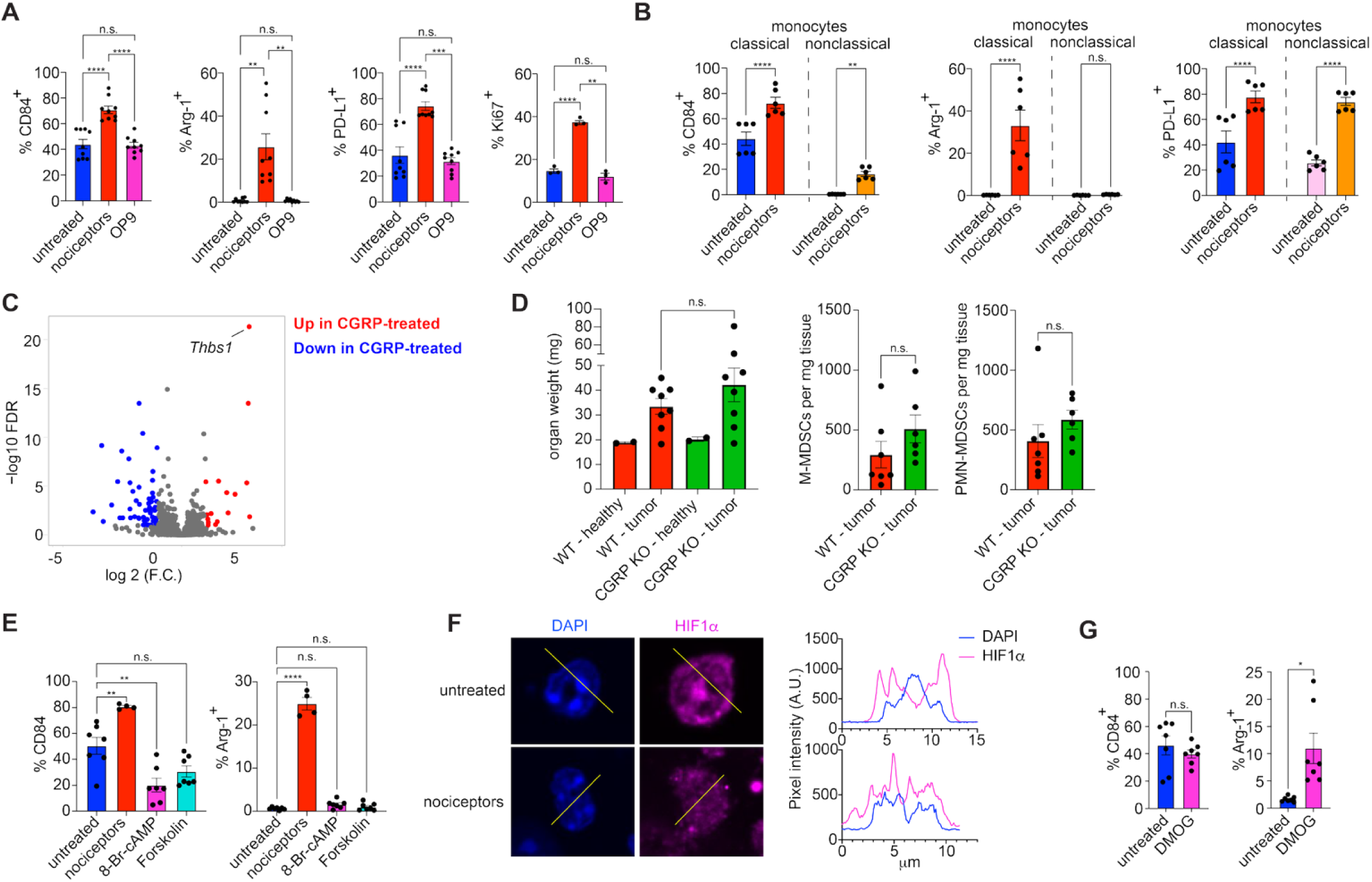
Nociceptors communicate with monocytes in a neuropeptide-independent manner. (**A**) FACS-purified Ly6C^Hi^ splenic monocytes were cultured alone or co-cultured with nociceptors or OP9 cells overnight, and the expression of indicated markers was assessed by flow cytometry. Mean +/- SEM of 3 independent experiments is shown. (**B**) FACS-purified Ly6C^Hi^ classical and Ly6C^Lo^ non-classical monocytes were co-cultured overnight with nociceptors, and the expression of indicated markers was assessed by flow cytometry. Mean +/-SEM of 2 independent experiments is shown. (**C**) FACS-purified Ly6C^Hi^ splenic monocytes were left untreated or treated with CGRP overnight and analyzed by RNAseq. (**D**) Bladders of CGRPα-deficient and control WT mice were implanted with MB49 cells. On day 14, organ weight as a proxy for tumor growth was measured, and MDSC accumulation was assessed by flow cytometry. Mean +/- SEM of 2 healthy and 8 tumor-implanted animals per group is shown. **(E)** FACS-purified Ly6C^Hi^ splenic monocytes were left untreated, cocultured with nociceptors or treated with 8-Br-cAMP (1mM) or Forskolin (50μM) overnight, and the expression of indicated markers was assessed by flow cytometry. Mean +/- SEM of 2 independent experiments is shown. (**F**) FACS-purified Ly6C^Hi^ splenic monocytes were cultured in the presence or absence of nociceptors, fixed, stained with DAPI and anti-Hif1α antibodies, and analyzed by confocal microscopy. (**G**) FACS-purified Ly6C^Hi^ splenic monocytes were treated with DMOG, and the expression of indicated markers was assessed by flow cytometry. Mean +/- SEM of 3 independent experiments is shown. One-way ANOVA with Tukey’s multiple comparisons test (**A,E**), two-way ANOVA with Tukey’s multiple comparisons test (**B**), and Welch’s t-test (**D**,**G**) were used for statistical analysis. *p<0.05, **p<0.01, ***p<0.001, ****p<0.0001

**Supplementary Figure 7.**
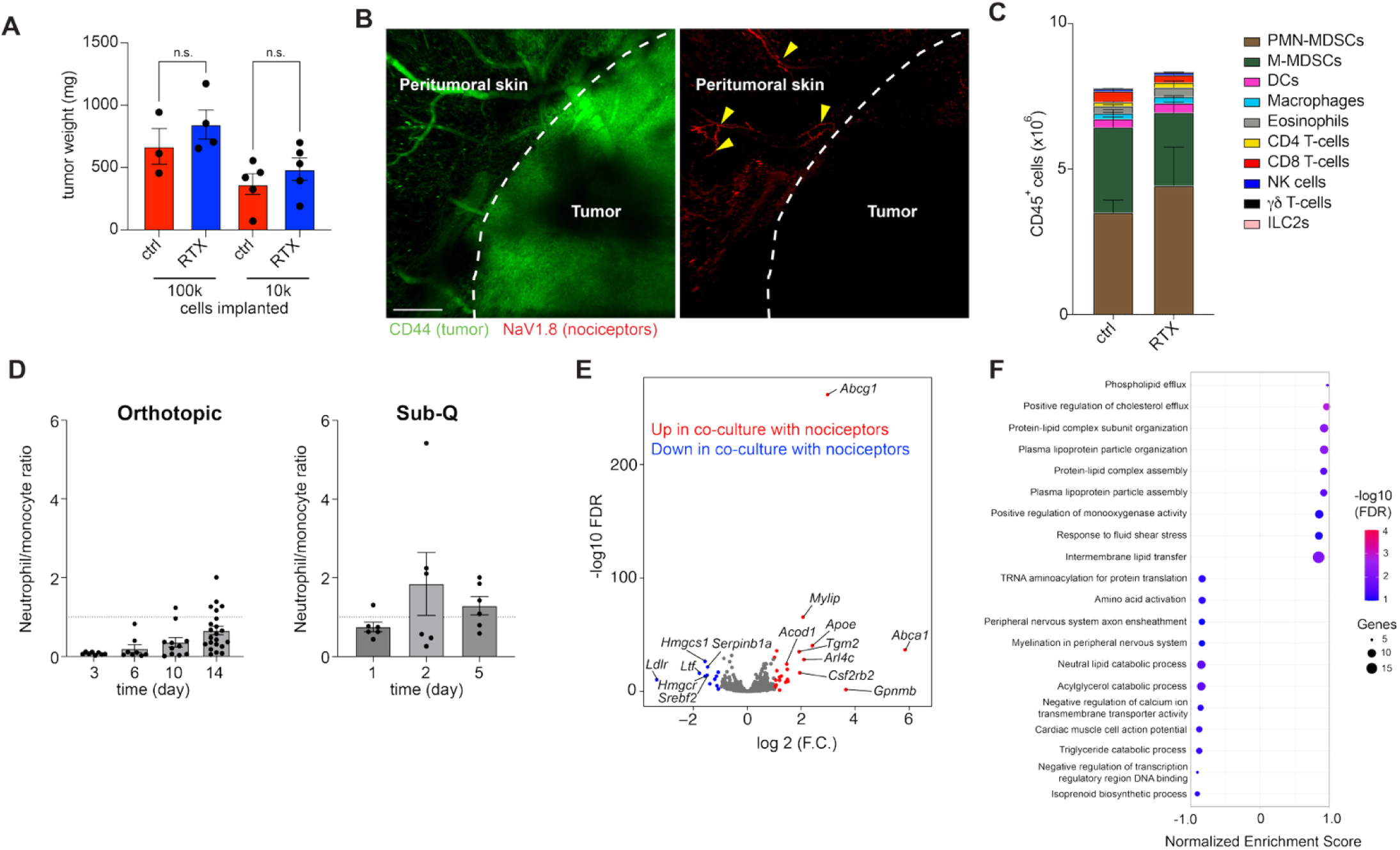
Nociceptors do not innervate subcutaneous MB49 tumors or induce MDSC phenotype in neutrophils. (**A**) Indicated number of MB49 cells were implanted sub-Q into nociceptor-ablated and -competent mice, and tumor weight was measured on day 14. Mean +/- SEM of 3-5 animals per group is shown. Welch’s t-test was used for statistical analysis. **(B)** MB49 cells were implanted sub-Q into nociceptor-reporter mice. 14 days later, the tumors were processed for iDISCO, stained for tumor-associated CD44, and imaged on a confocal microscope. Skin nociceptors are highlighted by arrowheads. (**C**) The abundance of indicated immune cells in D14 subcutaneous MB49 tumors was assessed by flow cytometry. Mean +/- SEM of 3 animals per group is shown. (**D**) Monocytic and neutrophilic infiltrates in orthotopic and sub-Q MB49 tumors were enumerated by flow cytometry at the indicated time points. Neutrophil to monocyte ratios are shown. Mean +/- SEM of 6-22 animals per group is shown. (**E**) Primary bone marrow neutrophils were cultured overnight alone or in the presence of nociceptors and analyzed by RNAseq. (**F**) GO terms enrichment analysis comparing the RNAseq datasets of neutrophils cultured with nociceptors or alone.

**Supplementary Figure 8.**
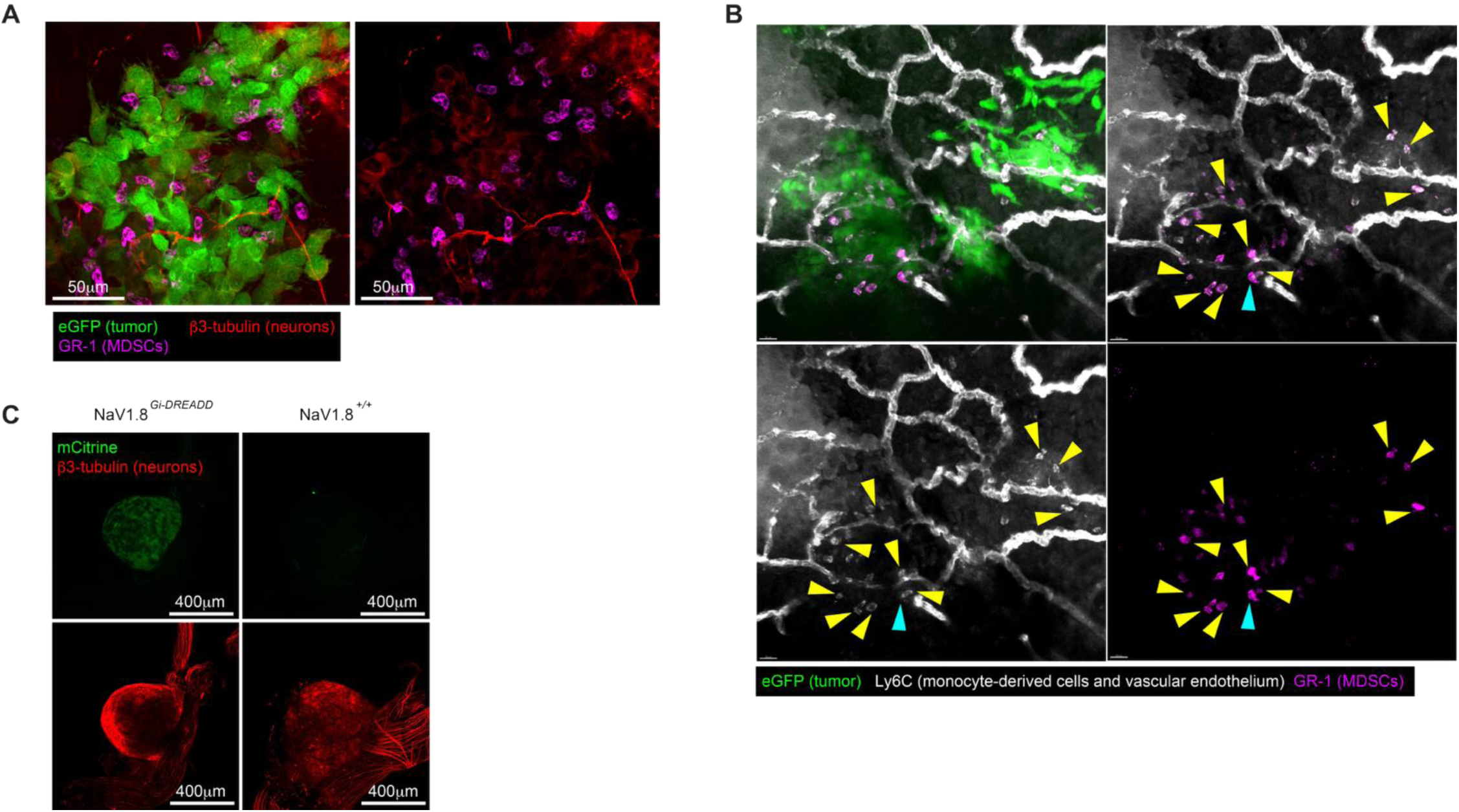
Tumor-infiltrating monocytic cells. (**A-B**) Bladders of eGFP-MB49-implanted mice were harvested on day 4, stained with the indicated antibodies, and analyzed as whole-mount preparations by confocal microscopy. (**B**) Monocyte-derived (GR-1^+^ Ly6C^+^) cells are highlighted by yellow, and neutrophils (GR-1^+^ Ly6C^dim^) by cyan arrows. (**C**) Dorsal root ganglia of NaV1.8*^Gi-DREADD^* and Cre-negative (NaV1.8^+/+^) littermates were stained with β3-tubulin antibodies and imaged on a confocal microscope

## SUPPLEMENTARY VIDEO LEGENDS

**Supplementary video 1. Nociceptors localize underneath the uroepithelium in normal bladder.** The urinary bladder of a nociceptor-reporter (NaV1.8^TdT^) mouse was fixed, split in half, stained with wheat germ agglutinin (WGA; cyan) and antibodies specific to E-cadherin (green) and laminin (white) to visualize the superficial layer, intermediate epithelium, and basement membrane, respectively. Stained bladders were imaged on a point-scanning confocal microscope. Nociceptors are shown in magenta.

**Supplementary video 2. Orthotopic MB49 tumors are innervated by nociceptors.** MB49 cells were implanted in the bladder of a nociceptor-reporter (NaV1.8^TdT^) mouse. The organ was harvested on day 14 and clarified for whole-mount imaging using the iDISCO protocol. Samples were stained with MAbs against CD44 and mCherry to visualize CD44^hi^ tumor cells and nociceptors, respectively. Imaging was performed on a point-scanning confocal microscope. Imaris® image analysis software was used to generate 3D reconstructions and to render the surface of the CD44^+^ tumor mass.

**Supplementary video 3. Day 2 MB49-implanted bladder.** The bladder of an eGFP-MB49-implanted mouse was harvested at day 1 post-implantation, stained with DAPI (blue) and antibodies against β3 tubulin (red) and GR-1 (magenta) to visualize nuclei, neurons, and myeloid cells of the monocyte/neutrophil lineage, respectively, and imaged as a whole-mount preparation on a point-scanning confocal microscope. GR-1^+^ MDSCs are highlighted by yellow arrows.

**Supplementary video 4. Day 4 MB49-implanted bladder.** The bladder of an eGFP-MB49-implanted mouse was harvested at day 4 post-implantation, stained with DAPI (blue) and antibodies against β3 tubulin (red) and GR-1 (magenta) to visualize nuclei, neurons, and myeloid cells of the monocyte/neutrophil lineage, respectively, and imaged as a whole-mount preparation on a point-scanning confocal microscope.

**Supplementary video 5. Day 7 MB49-implanted bladder.** The bladder of an eGFP-MB49-implanted mouse was harvested at day 7 post-implantation, stained with antibodies against β3 tubulin (red), GR-1 (magenta), and CD8 (white) to visualize neurons, myeloid cells of the monocyte/neutrophil lineage, and CD8 T-cells, respectively, and imaged as a whole-mount preparation on a point-scanning confocal microscope. CD8 T-cells are highlighted by yellow arrows.

## References

1 Deng, L., Gillis, J. E., Chiu, I. M. & Kaplan, D. H. Sensory neurons: An integrated component of innate immunity. Immunity 57, 815–831 (2024). 10.1016/j.immuni.2024.03.008

2 Hanč, P., Messou, M. A., Ajit, J. & von Andrian, U. H. Setting the tone: nociceptors as conductors of immune responses. Trends Immunol 45, 783–798 (2024). 10.1016/j.it.2024.08.007

3 de Visser, K. E. & Joyce, J. A. The evolving tumor microenvironment: From cancer initiation to metastatic outgrowth. Cancer Cell 41, 374–403 (2023). 10.1016/j.ccell.2023.02.016

4 Raskov, H., Orhan, A., Christensen, J. P. & Gogenur, I. Cytotoxic CD8(+) T cells in cancer and cancer immunotherapy. Br J Cancer 124, 359–367 (2021). 10.1038/s41416-020-01048-4

5 Burke, K. P., Chaudhri, A., Freeman, G. J. & Sharpe, A. H. The B7:CD28 family and friends: Unraveling coinhibitory interactions. Immunity 57, 223–244 (2024). 10.1016/j.immuni.2024.01.013

6 Cheng, X., Wang, H., Wang, Z., Zhu, B. & Long, H. Tumor-associated myeloid cells in cancer immunotherapy. J Hematol Oncol 16, 71 (2023). 10.1186/s13045-023-01473-x

7 Bronte, V. et al. Recommendations for myeloid-derived suppressor cell nomenclature and characterization standards. Nat Commun 7, 12150 (2016). 10.1038/ncomms12150

8 Li, X. et al. Targeting tumor innervation: premises, promises, and challenges. Cell Death Discov 8, 131 (2022). 10.1038/s41420-022-00930-9

9 Hanč, P. et al. Multimodal control of dendritic cell functions by nociceptors. Science 379, eabm5658 (2023). 10.1126/science.abm5658

10 Kashem, S. W. et al. Nociceptive Sensory Fibers Drive Interleukin-23 Production from CD301b+ Dermal Dendritic Cells and Drive Protective Cutaneous Immunity. Immunity 43, 515–526 (2015). 10.1016/j.immuni.2015.08.016

11 Riol-Blanco, L. et al. Nociceptive sensory neurons drive interleukin-23-mediated psoriasiform skin inflammation. Nature 510, 157–161 (2014). 10.1038/nature13199

12 Wu, M. et al. Innervation of nociceptor neurons in the spleen promotes germinal center responses and humoral immunity. Cell (2024). 10.1016/j.cell.2024.04.027

13 Baral, P. et al. Nociceptor sensory neurons suppress neutrophil and gammadelta T cell responses in bacterial lung infections and lethal pneumonia. Nat Med 24, 417–426 (2018). 10.1038/nm.4501

14 Chiu, I. M. et al. Bacteria activate sensory neurons that modulate pain and inflammation. Nature 501, 52-57 (2013). 10.1038/nature12479

15 Pinho-Ribeiro, F. A. et al. Bacteria hijack a meningeal neuroimmune axis to facilitate brain invasion. Nature 615, 472–481 (2023). 10.1038/s41586-023-05753-x

16 Lu, Y. Z. et al. CGRP sensory neurons promote tissue healing via neutrophils and macrophages. Nature 628, 604–611 (2024). 10.1038/s41586-024-07237-y

17 Mardelle, U., Bretaud, N., Daher, C. & Feuillet, V. From pain to tumor immunity: influence of peripheral sensory neurons in cancer. Front Immunol 15, 1335387 (2024). 10.3389/fimmu.2024.1335387

18 Vats, K. et al. Sensory Nerves Impede the Formation of Tertiary Lymphoid Structures and Development of Protective Antimelanoma Immune Responses. Cancer Immunol Res 10, 1141–1154 (2022). 10.1158/2326-6066.CIR-22-0110

19 Balood, M. et al. Nociceptor neurons affect cancer immunosurveillance. Nature 611, 405–412 (2022). 10.1038/s41586-022-05374-w

20 Darragh, L. B. et al. Sensory nerve release of CGRP increases tumor growth in HNSCC by suppressing TILs. Med 5, 254–270 e258 (2024). 10.1016/j.medj.2024.02.002

21 McIlvried, L. A., Atherton, M. A., Horan, N. L., Goch, T. N. & Scheff, N. N. Sensory Neurotransmitter Calcitonin Gene-Related Peptide Modulates Tumor Growth and Lymphocyte Infiltration in Oral Squamous Cell Carcinoma. Adv Biol (Weinh) 6, e2200019 (2022). 10.1002/adbi.202200019

22 Restaino, A. C. et al. Tumor-infiltrating nociceptor neurons promote immunosuppression. Sci Signal 18, eads7889 (2025). 10.1126/scisignal.ads7889

23 Zhi, X. et al. Nociceptive neurons promote gastric tumour progression via a CGRP-RAMP1 axis. Nature 640, 802–810 (2025). 10.1038/s41586-025-08591-1

24 Padmanaban, V. et al. Neuronal substance P drives metastasis through an extracellular RNA-TLR7 axis. Nature 633, 207–215 (2024). 10.1038/s41586-024-07767-5

25 Gyorffy, B. et al. An online survival analysis tool to rapidly assess the effect of 22,277 genes on breast cancer prognosis using microarray data of 1,809 patients. Breast Cancer Res Treat 123, 725–731 (2010). 10.1007/s10549-009-0674-9

26 Barker, P. A., Mantyh, P., Arendt-Nielsen, L., Viktrup, L. & Tive, L. Nerve Growth Factor Signaling and Its Contribution to Pain. J Pain Res 13, 1223–1241 (2020). 10.2147/JPR.S247472

27 Merighi, A. Brain-Derived Neurotrophic Factor, Nociception, and Pain. Biomolecules 14 (2024). 10.3390/biom14050539

28 Malin, S. A. et al. Glial cell line-derived neurotrophic factor family members sensitize nociceptors in vitro and produce thermal hyperalgesia in vivo. J Neurosci 26, 8588–8599 (2006). 10.1523/JNEUROSCI.1726-06.2006

29 White-Gilbertson, S., Davis, M., Voelkel-Johnson, C. & Kasman, L. M. Sex differences in the MB49 syngeneic, murine model of bladder cancer. Bladder (San Franc) 3 (2016). 10.14440/bladder.2016.73

30 Erman, A. et al. How cancer cells attach to urinary bladder epithelium in vivo: study of the early stages of tumorigenesis in an orthotopic mouse bladder tumor model. Histochem Cell Biol 151, 263–273 (2019). 10.1007/s00418-018-1738-x

31 Tham, S. M., Esuvaranathan, K. & Mahendran, R. A Murine Orthotopic Bladder Tumor Model and Tumor Detection System. J Vis Exp (2017). 10.3791/55078

32 Alshetaiwi, H. et al. Defining the emergence of myeloid-derived suppressor cells in breast cancer using single-cell transcriptomics. Sci Immunol 5 (2020). 10.1126/sciimmunol.aay6017

33 Raber, P., Ochoa, A. C. & Rodriguez, P. C. Metabolism of L-arginine by myeloid-derived suppressor cells in cancer: mechanisms of T cell suppression and therapeutic perspectives. Immunol Invest 41, 614–634 (2012). 10.3109/08820139.2012.680634

34 Ochando, J. C. & Chen, S. H. Myeloid-derived suppressor cells in transplantation and cancer. Immunol Res 54, 275–285 (2012). 10.1007/s12026-012-8335-1

35 Mack, M. et al. Expression and characterization of the chemokine receptors CCR2 and CCR5 in mice. J Immunol 166, 4697–4704 (2001). 10.4049/jimmunol.166.7.4697

36 Schneider, M. A. et al. In vitro and in vivo properties of a dimeric bispecific single-chain antibody IgG-fusion protein for depletion of CCR2+ target cells in mice. Eur J Immunol 35, 987–995 (2005). 10.1002/eji.200425512

37 Reich, B. et al. Fibrocytes develop outside the kidney but contribute to renal fibrosis in a mouse model. Kidney Int 84, 78–89 (2013). 10.1038/ki.2013.84

38 Huang, S. et al. Lymph nodes are innervated by a unique population of sensory neurons with immunomodulatory potential. Cell 184, 441–459 e425 (2021). 10.1016/j.cell.2020.11.028

39 Renier, N. et al. iDISCO: a simple, rapid method to immunolabel large tissue samples for volume imaging. Cell 159, 896–910 (2014). 10.1016/j.cell.2014.10.010

40 Olah, Z. et al. Ligand-induced dynamic membrane changes and cell deletion conferred by vanilloid receptor 1. J Biol Chem 276, 11021–11030 (2001). 10.1074/jbc.M008392200

41 Galon, J. & Bruni, D. Tumor Immunology and Tumor Evolution: Intertwined Histories. Immunity 52, 55–81 (2020). 10.1016/j.immuni.2019.12.018

42 Hutloff, A. et al. ICOS is an inducible T-cell co-stimulator structurally and functionally related to CD28. Nature 397, 263–266 (1999). 10.1038/16717

43 Hao, Z. & Rajewsky, K. Homeostasis of peripheral B cells in the absence of B cell influx from the bone marrow. J Exp Med 194, 1151–1164 (2001). 10.1084/jem.194.8.1151

44 Chen, W., Khilko, S., Fecondo, J., Margulies, D. H. & McCluskey, J. Determinant selection of major histocompatibility complex class I-restricted antigenic peptides is explained by class I-peptide affinity and is strongly influenced by nondominant anchor residues. J Exp Med 180, 1471–1483 (1994). 10.1084/jem.180.4.1471

45 Falk, K. et al. Both human and mouse cells expressing H-2Kb and ovalbumin process the same peptide, SIINFEKL. Cell Immunol 150, 447–452 (1993). 10.1006/cimm.1993.1212

46 Tiwari, A. et al. Towards a consensus definition of immune exclusion in cancer. Front Immunol 14, 1084887 (2023). 10.3389/fimmu.2023.1084887

47 Ombrato, L. et al. Metastatic-niche labelling reveals parenchymal cells with stem features. Nature 572, 603–608 (2019). 10.1038/s41586-019-1487-6

48 Ginhoux, F. & Jung, S. Monocytes and macrophages: developmental pathways and tissue homeostasis. Nat Rev Immunol 14, 392–404 (2014). 10.1038/nri3671

49 Hanč, P., Messou, M. A., Wang, Y. & von Andrian, U. H. Control of myeloid cell functions by nociceptors. Front Immunol 14, 1127571 (2023). 10.3389/fimmu.2023.1127571

50 Ashburner, M. et al. Gene ontology: tool for the unification of biology. The Gene Ontology Consortium. Nat Genet 25, 25–29 (2000). 10.1038/75556

51 Gene Ontology, C., et al. The Gene Ontology knowledgebase in 2023. Genetics 224 (2023). 10.1093/genetics/iyad031

52 Rodriguez, P. C., Ochoa, A. C. & Al-Khami, A. A. Arginine Metabolism in Myeloid Cells Shapes Innate and Adaptive Immunity. Front Immunol 8, 93 (2017). 10.3389/fimmu.2017.00093

53 Hegde, S., Leader, A. M. & Merad, M. MDSC: Markers, development, states, and unaddressed complexity. Immunity 54, 875–884 (2021). 10.1016/j.immuni.2021.04.004

54 Wang, H. Y. et al. Surface TREM2 on circulating M-MDSCs as a novel prognostic factor for adults with treatment-naive diffuse large B-cell lymphoma. Exp Hematol Oncol 12, 35 (2023). 10.1186/s40164-023-00399-x

55 Park, M. D. et al. TREM2 macrophages drive NK cell paucity and dysfunction in lung cancer. Nat Immunol 24, 792–801 (2023). 10.1038/s41590-023-01475-4

56 Gabrilovich, D. I. The Dawn of Myeloid-Derived Suppressor Cells: Identification of Arginase I as the Mechanism of Immune Suppression. Cancer Res 81, 3953–3955 (2021). 10.1158/0008-5472.CAN-21-1237

57 Garlanda, C. & Mantovani, A. Interleukin-1 in tumor progression, therapy, and prevention. Cancer Cell 39, 1023–1027 (2021). 10.1016/j.ccell.2021.04.011

58 Chun, E. et al. CCL2 Promotes Colorectal Carcinogenesis by Enhancing Polymorphonuclear Myeloid-Derived Suppressor Cell Population and Function. Cell Rep 12, 244–257 (2015). 10.1016/j.celrep.2015.06.024

59 Ren, X. et al. Inhibition of CCL7 derived from Mo-MDSCs prevents metastatic progression from latency in colorectal cancer. Cell Death Dis 12, 484 (2021). 10.1038/s41419-021-03698-5

60 Zhou, S. L. et al. CXCL5 contributes to tumor metastasis and recurrence of intrahepatic cholangiocarcinoma by recruiting infiltrative intratumoral neutrophils. Carcinogenesis 35, 597–605 (2014). 10.1093/carcin/bgt397

61 Zhang, Y. et al. CD39 inhibition and VISTA blockade may overcome radiotherapy resistance by targeting exhausted CD8+ T cells and immunosuppressive myeloid cells. Cell Rep Med 4, 101151 (2023). 10.1016/j.xcrm.2023.101151

62 Wang, J. C. & Sun, L. PD-1/PD-L1, MDSC Pathways, and Checkpoint Inhibitor Therapy in Ph(-) Myeloproliferative Neoplasm: A Review. Int J Mol Sci 23 (2022). 10.3390/ijms23105837

63 Mootha, V. K. et al. PGC-1alpha-responsive genes involved in oxidative phosphorylation are coordinately downregulated in human diabetes. Nat Genet 34, 267–273 (2003). 10.1038/ng1180

64 Subramanian, A. et al. Gene set enrichment analysis: a knowledge-based approach for interpreting genome-wide expression profiles. Proc Natl Acad Sci U S A 102, 15545–15550 (2005). 10.1073/pnas.0506580102

65 Patel, A. A. et al. The fate and lifespan of human monocyte subsets in steady state and systemic inflammation. J Exp Med 214, 1913–1923 (2017). 10.1084/jem.20170355

66 Jiang, X. X. et al. Optimization of seeding density of OP9 cells to improve hematopoietic differentiation efficiency. BMC Mol Cell Biol 25, 10 (2024). 10.1186/s12860-024-00503-x

67 Pinho-Ribeiro, F. A., Verri, W. A., Jr. & Chiu, I. M. Nociceptor Sensory Neuron-Immune Interactions in Pain and Inflammation. Trends Immunol 38, 5–19 (2017). 10.1016/j.it.2016.10.001

68 Wallrapp, A. et al. Calcitonin Gene-Related Peptide Negatively Regulates Alarmin-Driven Type 2 Innate Lymphoid Cell Responses. Immunity 51, 709–723 e706 (2019). 10.1016/j.immuni.2019.09.005

69 Carter, M. E., Soden, M. E., Zweifel, L. S. & Palmiter, R. D. Genetic identification of a neural circuit that suppresses appetite. Nature 503, 111–114 (2013). 10.1038/nature12596

70 Han, H. et al. TRRUST v2: an expanded reference database of human and mouse transcriptional regulatory interactions. Nucleic Acids Res 46, D380–D386 (2018). 10.1093/nar/gkx1013

71 Kimura, K., Jackson, T. L. B. & Huang, R. C. C. Interaction and Collaboration of SP1, HIF-1, and MYC in Regulating the Expression of Cancer-Related Genes to Further Enhance Anticancer Drug Development. Curr Issues Mol Biol 45, 9262–9283 (2023). 10.3390/cimb45110580

72 Colegio, O. R. et al. Functional polarization of tumour-associated macrophages by tumour-derived lactic acid. Nature 513, 559–563 (2014). 10.1038/nature13490

73 Corzo, C. A. et al. HIF-1alpha regulates function and differentiation of myeloid-derived suppressor cells in the tumor microenvironment. J Exp Med 207, 2439–2453 (2010). 10.1084/jem.20100587

74 Taylor, C. T. & Scholz, C. C. The effect of HIF on metabolism and immunity. Nat Rev Nephrol 18, 573–587 (2022). 10.1038/s41581-022-00587-8

75 Ayrapetov, M. K. et al. Activation of Hif1alpha by the prolylhydroxylase inhibitor dimethyoxalyglycine decreases radiosensitivity. PLoS One 6, e26064 (2011). 10.1371/journal.pone.0026064

76 Giese, M. A., Hind, L. E. & Huttenlocher, A. Neutrophil plasticity in the tumor microenvironment. Blood 133, 2159–2167 (2019). 10.1182/blood-2018-11-844548

77 Zhu, H. et al. Cre-dependent DREADD (Designer Receptors Exclusively Activated by Designer Drugs) mice. Genesis 54, 439–446 (2016). 10.1002/dvg.22949

78 Roth, B. L. DREADDs for Neuroscientists. Neuron 89, 683–694 (2016). 10.1016/j.neuron.2016.01.040

79 von Andrian, U. H. & Mackay, C. R. T-cell function and migration. Two sides of the same coin. N Engl J Med 343, 1020–1034 (2000). 10.1056/NEJM200010053431407

80 Saluja, M. & Gilling, P. Intravesical bacillus Calmette-Guerin instillation in non-muscle-invasive bladder cancer: A review. Int J Urol 25, 18–24 (2018). 10.1111/iju.13410

81 Teoh, J. Y. et al. Recurrence mechanisms of non-muscle-invasive bladder cancer - a clinical perspective. Nat Rev Urol 19, 280–294 (2022). 10.1038/s41585-022-00578-1

82 Klein, C. et al. Myeloid-Derived Suppressor Cells in Bladder Cancer: An Emerging Target. Cells 13 (2024). 10.3390/cells13211779

83 Bazargan, S. et al. Targeting myeloid-derived suppressor cells with gemcitabine to enhance efficacy of adoptive cell therapy in bladder cancer. Front Immunol 14, 1275375 (2023). 10.3389/fimmu.2023.1275375

84 Takeyama, Y. et al. Myeloid-derived suppressor cells are essential partners for immune checkpoint inhibitors in the treatment of cisplatin-resistant bladder cancer. Cancer Lett 479, 89–99 (2020). 10.1016/j.canlet.2020.03.013

85 Yang, G. et al. Accumulation of myeloid-derived suppressor cells (MDSCs) induced by low levels of IL-6 correlates with poor prognosis in bladder cancer. Oncotarget 8, 38378–38388 (2017). 10.18632/oncotarget.16386

86 Hafner, C. et al. Evidence for oligoclonality and tumor spread by intraluminal seeding in multifocal urothelial carcinomas of the upper and lower urinary tract. Oncogene 20, 4910–4915 (2001). 10.1038/sj.onc.1204671

87 Bohle, A. et al. Inhibition of bladder carcinoma cell adhesion by oligopeptide combinations in vitro and in vivo. J Urol 167, 357–363 (2002).

88 Yang, H. et al. HMGB1 released from nociceptors mediates inflammation. Proc Natl Acad Sci U S A 118 (2021). 10.1073/pnas.2102034118

89 Simeoli, R. et al. Exosomal cargo including microRNA regulates sensory neuron to macrophage communication after nerve trauma. Nat Commun 8, 1778 (2017). 10.1038/s41467-017-01841-5

90 Liu, Y. R., Yin, P. N., Silvers, C. R. & Lee, Y. F. Enhanced metastatic potential in the MB49 urothelial carcinoma model. Sci Rep 9, 7425 (2019). 10.1038/s41598-019-43641-5

91 Mani, K. et al. Causes of death among people living with metastatic cancer. Nat Commun 15, 1519 (2024). 10.1038/s41467-024-45307-x

92 Beigi, A., Vafaei-Nodeh, S., Huang, L., Sun, S. Z. & Ko, J. J. Survival Outcomes Associated with First and Second-Line Palliative Systemic Therapies in Patients with Metastatic Bladder Cancer. Curr Oncol 28, 3812–3824 (2021). 10.3390/curroncol28050325

93 Shinagare, A. B. et al. Metastatic pattern of bladder cancer: correlation with the characteristics of the primary tumor. AJR Am J Roentgenol 196, 117–122 (2011). 10.2214/AJR.10.5036

94 Dyrskjot, L. et al. Bladder cancer. Nat Rev Dis Primers 9, 58 (2023). 10.1038/s41572-023-00468-9

95 Miller, A., Burson, H., Soling, A. & Roughan, J. Welfare Assessment following Heterotopic or Orthotopic Inoculation of Bladder Cancer in C57BL/6 Mice. PLoS One 11, e0158390 (2016). 10.1371/journal.pone.0158390

96 Yosipovitch, G. et al. Association of pain and itch with depth of invasion and inflammatory cell constitution in skin cancer: results of a large clinicopathologic study. JAMA Dermatol 150, 1160–1166 (2014). 10.1001/jamadermatol.2014.895

97 Hanes, W. M. et al. Neuronal Circuits Modulate Antigen Flow Through Lymph Nodes. Bioelectron Med 3, 18–28 (2016). 10.15424/bioelectronmed.2016.00001

98 Nassar, M. A. et al. Nociceptor-specific gene deletion reveals a major role for Nav1.7 (PN1) in acute and inflammatory pain. Proc Natl Acad Sci U S A 101, 12706–12711 (2004). 10.1073/pnas.0404915101

99 Erman, A. et al. How Cancer Cells Invade Bladder Epithelium and Form Tumors: The Mouse Bladder Tumor Model as a Model of Tumor Recurrence in Patients. Int J Mol Sci 22 (2021). 10.3390/ijms22126328

100 Zhao, L. et al. Exploration of CRISPR/Cas9-based gene editing as therapy for osteoarthritis. Ann Rheum Dis 78, 676–682 (2019). 10.1136/annrheumdis-2018-214724

101 Hanč, P. & von Andrian, U. H. In vitro culture of adult mouse DRG neurons. Bio-protocol Preprint bio-protocol.org/prep2876. (2025).

102 Picelli, S. et al. Smart-seq2 for sensitive full-length transcriptome profiling in single cells. Nat Methods 10, 1096–1098 (2013). 10.1038/nmeth.2639

103 Picelli, S. et al. Full-length RNA-seq from single cells using Smart-seq2. Nat Protoc 9, 171–181 (2014). 10.1038/nprot.2014.006

104 Ge, S. X., Son, E. W. & Yao, R. iDEP: an integrated web application for differential expression and pathway analysis of RNA-Seq data. BMC Bioinformatics 19, 534 (2018). 10.1186/s12859-018-2486-6

105 Zhou, Y. et al. Metascape provides a biologist-oriented resource for the analysis of systems-level datasets. Nat Commun 10, 1523 (2019). 10.1038/s41467-019-09234-6

106 Schindelin, J., et al. Fiji: an open-source platform for biological-image analysis. Nat Methods 9, 676–682 (2012). 10.1038/nmeth.2019

107 Bankhead, P. et al. QuPath: Open source software for digital pathology image analysis. Sci Rep 7, 16878 (2017). 10.1038/s41598-017-17204-5

108 Arganda-Carreras, I. et al. Trainable Weka Segmentation: a machine learning tool for microscopy pixel classification. Bioinformatics 33, 2424–2426 (2017). 10.1093/bioinformatics/btx180

109 Simunic, I., Jagecic, D., Isakovic, J., Dobrivojevic Radmilovic, M. & Mitrecic, D. Lusca: FIJI (ImageJ) based tool for automated morphological analysis of cellular and subcellular structures. Sci Rep 14, 7383 (2024). 10.1038/s41598-024-57650-6

110 Gyorffy, B. Integrated analysis of public datasets for the discovery and validation of survival-associated genes in solid tumors. Innovation (Camb) 5, 100625 (2024). 10.1016/j.xinn.2024.100625

